# Cholecystokinin-A Signaling Regulates Automaticity of Pacemaker Cardiomyocytes

**DOI:** 10.1101/2023.01.24.525392

**Authors:** Hongmei Ruan, Ravi Mandla, Namita Ravi, Giselle Galang, Amanda W. Soe, Jeffrey E. Olgin, Di Lang, Vasanth Vedantham

**Author notes:** <u>Corresponding Authors:</u> Hongmei Ruan, MD, PhD, Smith Cardiovascular Research Building, 555 Mission Bay Blvd South, 314, San Francisco, CA 94158, Vasanth Vedantham, MD, PhD, Smith Cardiovascular Research Building, 555 Mission Bay Blvd South, 352M, San Francisco, CA 94158. These authors contributed equally to this work and share senior authorship (Hongmei Ruan, Vasanth Vedantham). Department of Medicine, Brigham and Women’s Hospital, Boston MA.

## Abstract

**Aims:** The behavior of pacemaker cardiomyocytes (PCs) in the sinoatrial node (SAN) is modulated by neurohormonal and paracrine factors, many of which signal through G-protein coupled receptors (GPCRs). The aims of the present study are to catalog GPCRs that are differentially expressed in the mammalian SAN and to define the acute physiological consequences of activating the cholecystokinin-A signaling system in isolated PCs.

**Methods and Results:** Using bulk and single cell RNA sequencing datasets, we identify a set of GPCRs that are differentially expressed between SAN and right atrial tissue, including several whose roles in PCs and in the SAN have not been thoroughly characterized. Focusing on one such GPCR, Cholecystokinin-A receptor (CCK_A_R), we demonstrate expression of *Cckar* mRNA specifically in mouse PCs, and further demonstrate that subsets of SAN fibroblasts and neurons within the cardiac intrinsic nervous system express cholecystokinin, the ligand for CCK_A_R. Using mouse models, we find that while baseline SAN function is not dramatically affected by loss of CCK_A_R, the firing rate of individual PCs is slowed by exposure to sulfated cholecystokinin-8 (sCCK-8), the high affinity ligand for CCK_A_R. The effect of sCCK-8 on firing rate is mediated by reduction in the rate of spontaneous phase 4 depolarization of PCs and is mitigated by activation of beta-adrenergic signaling.

**Conclusions:** (1) PCs express many GPCRs whose specific roles in SAN function have not been characterized, (2) Activation of the the cholecystokinin-A signaling pathway regulates PC automaticity.

## INTRODUCTION

The sinoatrial node (SAN) contains an electrotonically coupled network of pacemaker cardiomyocytes (PCs) that depolarizes the atrial myocardium to initiate the heartbeat. Cellular automaticity in PCs is driven by mutual entrainment of hyperpolarization-activated cyclic nucleotide gated ion channels (“membrane clock”) and spontaneous subsarcolemmal calcium release (“calcium clock”) ^1^. PCs are also densely innervated with sympathetic ^2^ and post-ganglionic parasympathetic nerve fibers ^3^, the latter originating within the ganglia of the cardiac intrinsic nervous system (CINS). In addition, paracrine signals from fibroblasts and macrophages, tissue stretch, and humoral input all influence the firing rate of PCs and impulse transmission from the SAN to the atrium. Incorporation of PCs and their complex responses into a patterned network allows the SAN to function as a robust and tunable rhythm generator for the heart.

In this context, it is noteworthy that the complexity of interactions between PCs and their various inputs are only beginning to be unraveled. While the model of mutual antagonism between sympathetic input via norepinephrine and parasympathetic input via acetylcholine explains much SAN behavior, other neurotransmitters and modulators, operating both within the CINS as well as at the CINS-SAN interface, are relevant to SAN function ^4^. Supporting the notion of additional pathways relevant to heart rate regulation, recent genome-wide expression studies by our group and other groups have highlighted the enrichment of mRNA encoding several neurotransmitters and neurotransmitter receptors within the CINS and SAN whose specific roles are not well understood, in addition to numerous G-protein coupled receptors (GPCRs) that are differentially expressed in PCs as compared to atrial cardiomyocytes. While physiological roles and functional consequences have been assigned to some of these pathways, many remain uncharacterized.

Here, we survey the landscape of differentially expressed GPCRs in mouse PCs and explore the cholecystokinin signaling pathway, which has not been well characterized in the SAN. In numerous organs and tissues, including the brain and the peripheral nervous system, activation of cholecystokinin A receptor (CCK_A_R) by sulfated cholecystokinin octapeptide (sCCK-8), its highest affinity agonist, results in signaling via the heterotrimeric G proteins (typically Gq/11 or Gs) with diverse downstream effects, including induction of gene expression, enzyme secretion, promotion of satiety, natriuresis, and a host of other important physiological consequences ^5^. However, much less is known about potential roles for CCK_A_ signaling in the SAN although its components have been identified in the heart from early stages of cardiac development ^6^. One prior study found that administration of sCCK-8 to isolated rat hearts results in heart rate slowing that persists despite autonomic blockade ^7^, while a separate study found that low-dose sCCK-8 led to tachycardia in rat hearts while high dose sCCK-8 induced bradycardia ^8^, suggesting a potentially complex mechanism of action on heart rhythm. However, the possibility of a specific chronotropic effect of cholecystokinin on PCs has not yet been demonstrated, nor has it been determined how activation of this pathway might be relevant to impulse generation and propagation within the isolated SAN.

In the present work, we mine existing mRNA expression datasets to demonstrate that *Cckar* is among the most differentially expressed GPCRs between murine SAN and right atrial cardiomyocytes ^9^. We further explore scRNA-seq datasets to define the specific SAN cell populations that express *Cckar* and its ligand, CCK. We then use cellular electrophysiology and optical mapping of SAN tissue in WT and *Cckar*^*-/-*^ mice to assess acute physiological responses to activation of the CCK_A_R signaling pathway in the mouse heart. Taken together, our data demonstrate a novel signaling axis between the CINS, SAN fibroblasts, and PCs that can regulate cardiac automaticity.

## 1. METHODS

A detailed methods section is available in the Supplementary Materials.

### 2.1 Animal Handling and Maintenance

Mouse studies were performed under IACUC-approved protocols at the University of California, San Francisco. In accordance with our protocol, animals were euthanized either by carbon dioxide asphyxiation followed by cervical dislocation, or by induction of deep anesthesia with 5% isoflurane, followed by surgical removal of the heart. *Cckar*^*tm1Kpn*^ (hereafter, *Cckar*^*-/-*^) mice were generated by Alan Kopin et al. and were obtained from Nirao Shah’s lab ^10^. *Cck*^*tm1*.*1(cre)Zjh*^ (hereafter, *Cck*^*Cre*^) mice were obtained from Jackson Labs. *Gt(ROSA)26Sor*^*tm4(ACTB-tdTomato*,*-EGFP)Luo*^ (hereafter, *ROSA*^*mT/mG*^) mice were generated by Muzumdar et al. and were obtained from Brian Black’s lab ^11^.

### 2.2 Analysis of Gene Expression

Bulk RNA sequencing data was generated by Vedantham et al. ^9^ and Galang et al. ^12^ and downloaded from the Gene Expression Omnibus (GEO, GSE65658 and GSE148515, respectively), while mouse E16.5 SAN single-cell RNA-sequencing data were generated by Goodyer et al. ^13^ and downloaded from GEO (GSE132658). Adult rabbit single-cell RNA-seq data were generated by Liang et al. ^14^ and were downloaded as fastq files from the NCBI Sequence Read Archive (PRJNA531288). Quantitative PCR was performed using Taqman Probes (Thermo Fisher) with GAPDH as an endogenous control using the ΔΔCT method for comparison of SAN and right atrial tissue. Immunostaining and acetylcholinesterase staining were performed as previously described and are detailed in the Supplementary Methods.

### 2.3 Transmitter Implantation and Analysis of Heart Rhythm Data

Adult mice (8-12 weeks old) underwent subcutaneous implantation of telemetry devices (ETA-F10, Data Science International, MN) using 2% inhaled isoflurane for anesthesia according to the manufacturer’s instructions. Heart rhythm analysis was performed offline using Ponemah (Data Sciences International, St Paul, MN). 48 hours of heart rhythm data were analyzed for each mouse. Time-domain, frequency-domain, and arrhythmia analysis were performed as previously described^15^ with modifications detailed in the Supplementary Methods.

### 2.4 Whole Cell Patch Clamp

Adult mouse pacemaker cells were isolated for recording as previously described ^12^ and as detailed in the Supplementary Methods. Action potentials were acquired from spontaneously beating adult cells of the indicated genotypes via the perforated patch-clamp technique with an Axopatch-700B amplifier using pCLAMP10.3. The pCLAMP 10.6 Clampfit module and LabChart7 were used for analysis of action potentials (APs) as detailed in the Supplementary Methods.

## 3 RESULTS

### 3.1 Differential GPCR Expression in the Sinoatrial Node

To identify GPCRs that might regulate mouse SAN function, we analyzed a previously published bulk RNA sequencing dataset generated by our lab that was derived from sorted neonatal mouse PCs and right atrial cardiomyocytes (RACMs) ^12^. Differentially expressed GPCRs were identified by intersecting the list of expressed genes in both PCs and RACMs with an annotated list of GPCRs obtained from the International Union of Basic and Clinical Pharmacology^16^. We identified a total of 71 GPCRs expressed in PCs or RACMs, of which 29 were enriched in RACMs, 22 enriched in the PCs, and 29 expressed in both based on a false discovery rate of 0.2 (Supplementary Figure 1, Supplementary Table). Among the list of GPCRs enriched in PCs were receptors for canonical regulators of SAN function including adenosine (*Adora1*) and norepinephrine (*Adra1b*), as well as enrichment of receptors with ligands that do not have established associations with SAN function, such as lysophosphatidic acid (*Lpar3*) and phoenixin (*Gpr173*), orphan GPCRs (*Gpr22, Gpr85, Gpr157*, and *Gpr27*), adhesion GPCRs (*Adgrb2, Adgrd1*), and an opsin (Opn3). A third category of GPCRs included those whose ligands are known to affect SAN function but via mechanisms that have not been fully characterized. This group includes receptors for ligands that are reported to acutely increase heart rate such as glucagon (*Gcgr*) ^17^, pituitary adenylate cyclase activating polypeptide (*Adcyap1r1*) ^18^, prostaglandin F_2α_ (*Ptgfr*) ^19^, and thromboxane A2 (*Tbx2ar*) ^20^; receptors whose ligands are reported to acutely decrease heart rate including sphingosine 1-phosphate (*S1pr3*) ^21^ and somatostatin (*Sstr4*) ^22^, and one ligand for highly expressed GPCR for which there exists prior conflicting data on acute physiological responses, cholecystokinin (*Cckar*). Given the unclear effects of cholecystokinin-A signaling in the PCs, and its high degree of differential expression with RACMs, we sought to define the role of this signaling pathway in more detail.

### 3.2 Cckar and Other GPCRs are Expressed in Sinoatrial Node Pacemaker Cells

To define the expression pattern of *Cckar* in the SAN at different time points, we explored publicly available RNAseq datasets of adult and embryonic SAN tissue. When the SAN was micro-dissected using laser capture, *Cckar* mRNA was highly enriched in SAN compared to right atrial tissue at embryonic day (E)14.5, postnatal day (P)4, and at P14 (Figure 1A-B). Additional RNA sequencing datasets derived from sorted cells enriched for pacemaker cells also confirmed expression of *Cckar* in late embryonic and neonatal murine SAN (Figure 1B). To test whether *Cckar* is expressed in adult SAN tissue, SAN tissue and right atrial tissue were dissected from 3 8-week-old adult hearts and *Cckar* mRNA was quantified using GAPDH as an endogenous control. *Cckar* mRNA was detected in all SAN samples but in none of the RA samples (Figure 1C).

**Figure 1.**
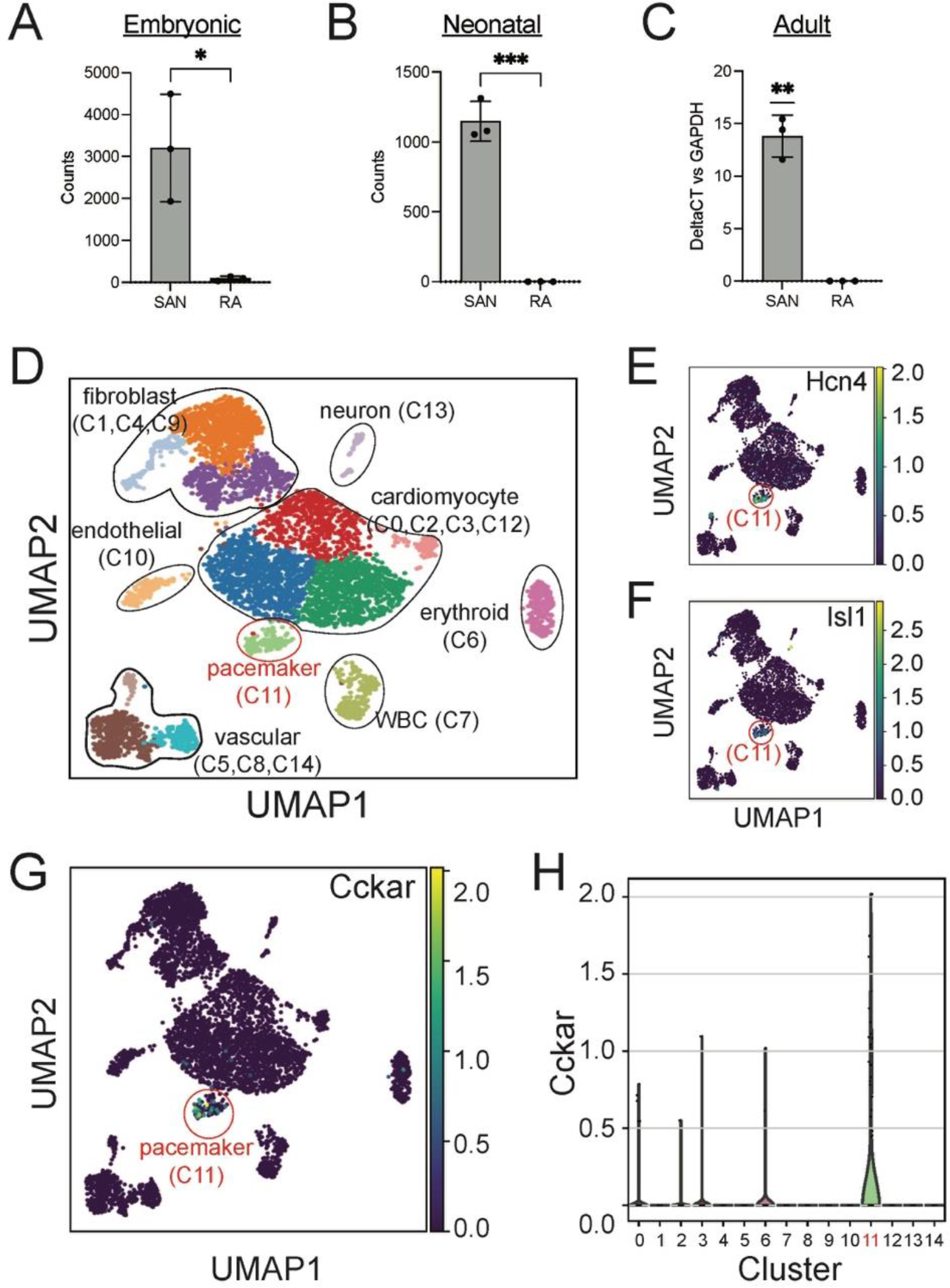
Cckar is Expressed in Mouse Cardiac Pacemaker Cells. (A) Read counts for Cckar from Bulk RNA sequencing of sinoatrial node (SAN) tissue and right atrial tissue isolated from 3 embryonic day 14.5 Hcn4-GFP BAC transgenic mouse embryos using laser capture microdissection (original dataset from Ref. [9]) A two-tailed t-test was used to assess for significance. ‘*’ denotes P < 0.05 (B) Read counts for *Cckar* from Bulk RNA sequencing of sorted Hcn4-GFP+ pacemaker cells and atrial cardiomyocytes from neonatal mouse hearts shown in the heatmap. A two-tailed t-test was used to assess for significance. ‘***’ denotes P < 0.001. (C) quantitative PCR for *Cckar* mRNA in samples manually dissected from adult mouse SAN and right atrium, using GAPDH as an endogenous control (*n* = 3 biological replicates). A one-sample t-test was used to assess for significance since *Cckar* was not detected in the RA samples. ‘**’ denotes P < 0.01 (D) Uniform Manifold Approximation Projection (UMAP) plot of leiden clustering analysis of single cell RNA sequencing data derived from 6 E16.5 mouse embryonic heart tissue samples taken from the SAN region demonstrated a pacemaker cell population (red circle, cluster 11), marked by co-expression of *Hcn4* (E) and *Isl1* (F), as well as several other identifiable cell types (original dataset from ref [12]). Cckar was differentially expressed in the pacemaker cluster (C11), as seen in the UMAP plot (G) and in the violin plot (H).

The SAN contains pacemaker cells, atrial cardiomyocytes, fibroblasts, and other cell types, so to establish whether *Cckar* expression is restricted to pacemaker cells or is also expressed in other SAN cell populations, we explored recently published single cell RNA sequencing datasets derived from murine and rabbit sinoatrial node. Datasets were clustered using UMAP and cell populations identified by expression of established marker genes (Figure 1D). A pacemaker cell cluster was identified that co-expressed *Hcn4* and *Isl1* (Figure 1E, F). Examination of *Cckar* expression within the *Hcn4*^*+*^*/Isl1*^*+*^ pacemaker cell cluster revealed high and differential expression as compared to other cell types. (Figure 1G, H and Supplementary Figure 2), confirming the presence of *Cckar* transcript specifically in mouse and rabbit cardiac pacemaker cells.

### 3.3 CCK-expressing neurons and fibroblasts are present in the SAN

Because sCCK-8 is produced by many cell types and can function via endocrine signaling pathways as well as via neurotransmission, we hypothesized that sCCK-8 might be produced within the cells of the cardiac intrinsic nervous system that project to the SAN or in other cardiac cell types near pacemaker cells. The scRNA-seq datasets described above demonstrated that *Cck* transcript was observed in both SAN fibroblasts and within a subset of cardiac neurons lying within the SAN (Figure 2A, B). To test further whether we could observe evidence of *Cck* expression in these cell types, we crossed *Cck*^*cre*^ mice, in which *Cre recombinase* is inserted downstream of the endogenous *Cck* gene using an internal ribosomal entry site, with *ROSA*^*mTmG*^ reporter mice, in which membrane targeted GFP is expressed upon recombination by Cre. Thus, GFP^+^ cells in *CCK-cre; ROSA*^*mTmG*^ represent cells that have expressed *Cck* for sufficient duration and in sufficient quantity to recombine a reporter allele at the *ROSA* locus.

**Figure 2.**
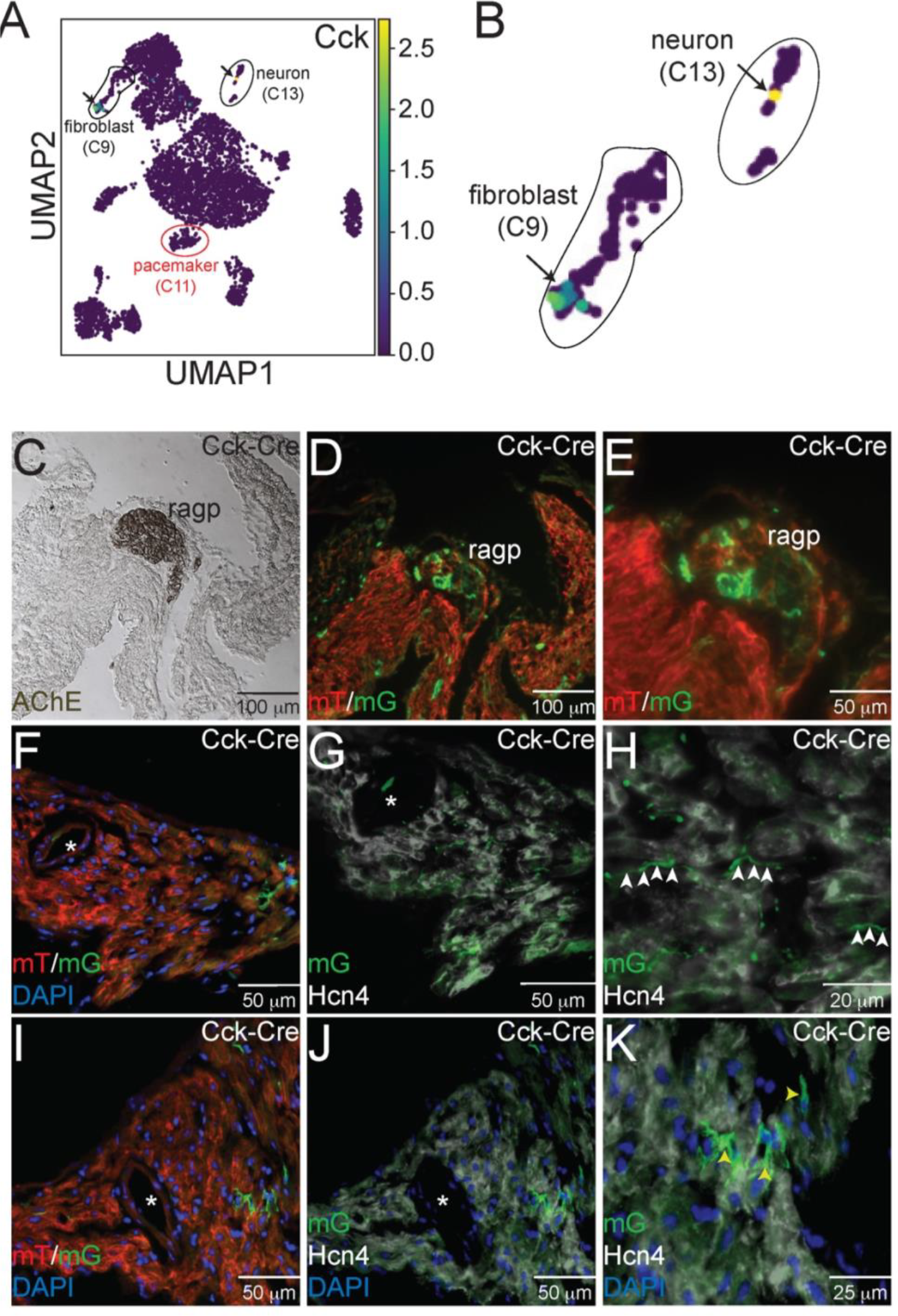
Expression of Cck in Neurons and Cardiac Fibroblasts. (A) Uniform Manifold Approximation and Projection (UMAP) plot of a single cell RNA sequencing dataset from neonatal sinoatrial node (SAN) (Ref [12]) demonstrates expression of *Cck* in a population of fibroblasts and in a cardiac neuron, as seen in the magnified images in panel (B). Adjacent serial sections from a neonatal *Cck-Cre;ROSA*^*mTmG*^ mouse heart stained with acetylcholinesterase to mark the right atrial cardiac ganglion (C) or imaged with epifluorescence to visualize Td-tomato and GFP expression (D,E) demonstrate that GFP+ cells, representing Cck expressing cells, are detectable in neuronal cell bodies within the ganglia. Sectioning and epifluorescence visualization of the SAN *Cck-Cre;ROSA*^*mTmG*^ showed few GFP+ cells (F) but as shown in (G) and (H), GFP+ axons could be observed coursing alongside Hcn4+ pacemaker cells (white arrowheads). Notably, GFP expression did not overlap with Hcn4 expression. A nearby section from the SAN (I) also revealed GFP+ cells with fibroblast morphologies in the SAN that did not express Hcn4 (J, K – yellow arrowheads).

Consecutive frozen sections from 1 month old *Cck*^*cre*^; *ROSA*^*mTmG*^ hearts were visualized under direct fluorescence and stained with acetyl cholinesterase (AChE), a marker for ganglionated plexi (Figure 2C). Adjacent sections demonstrated GFP+ neurons within the right atrial ganglionated plexus (Figure 2D, E). In addition, GFP+ cellular projections were observed to course within the SAN but did not overlap with Hcn4 expression, a marker of cardiac pacemaker cells (Figure 2F-H). The morphology and locations of these projections suggest neuronal identity. In addition to these projections, we also observed several Hcn4^-^ non-myocyte cells that expressed GFP, with morphologies consistent with fibroblasts (Figure 2I-K). Taken together, the imaging and scRNA-seq data support the conclusions that CCK-producing neurons are present within the heart and project to the SAN and that the SAN contains CCK-expressing fibroblasts. Importantly, these data establish that either the CINS or native SAN fibroblasts (or both) could serve as sources of ligand for Cckar expressed on cardiac pacemaker cells.

### 3.4 Sinoatrial Node Function in *Cckar*^*-/-*^ Mice

To test whether CCK_A_R is required for SAN function, we assessed heart rhythm and activity using implanted transmitters in *Cckar*^*-/-*^ mice (*n*=11, 8 male and 3 female) in comparison with WT littermates (n=7, 5 male and 2 female). EKG tracings from both groups of mice were similar with no significant differences in average PR or QRS intervals (Figure 3A, Supplementary Figure 3A-E). With respect to heart rate, the *Cckar*^*-/-*^ mice exhibited a modest slowing of average heart rate, particularly during waking hours (high activity period) (Figure 3B). Notably, activity counts were also reduced in the *Cckar*^*-/-*^ mice during similar periods. (Figure 3C). Both intrinsic heart rate (determined after administration of atropine and propranolol under anesthesia) and maximum heart rate (determined after isoproterenol infusion in awake mice) were similar between *Cckar*^*-/-*^ mice and WT littermates (Figure 3D, E). When heart rate was collated into 10 beat-per-minute bins and plotted in a histogram, there was a modest shift to slightly slower heart rates during high activity periods in the knockout mice, in keeping with their reduced activity counts during these periods. (Figure 3F). For each mouse, we also counted sinus pauses, episodes of sinus bradycardia, and episodes of AV block over 48 hours of continuous recording and found no significant differences in arrhythmia frequency between WT and *Cckar*^*-/-*^ mice (Supplementary Figure 3F-H). We also assessed heart rate variability in *Cckar*^*-/-*^ mice in the time domain by estimating RMSSD, SDRR, and pNN6 during low activity periods, and did not observe any significant differences between the two groups of mice (Supplementary Figure 4A-F). Frequency domain analysis revealed a small increase in low frequency power in *Cckar*^*-/-*^ mice without significant differences in total power or in high and frequency contributions to the power spectra (Supplementary Figure 4G-J).

**Figure 3.**
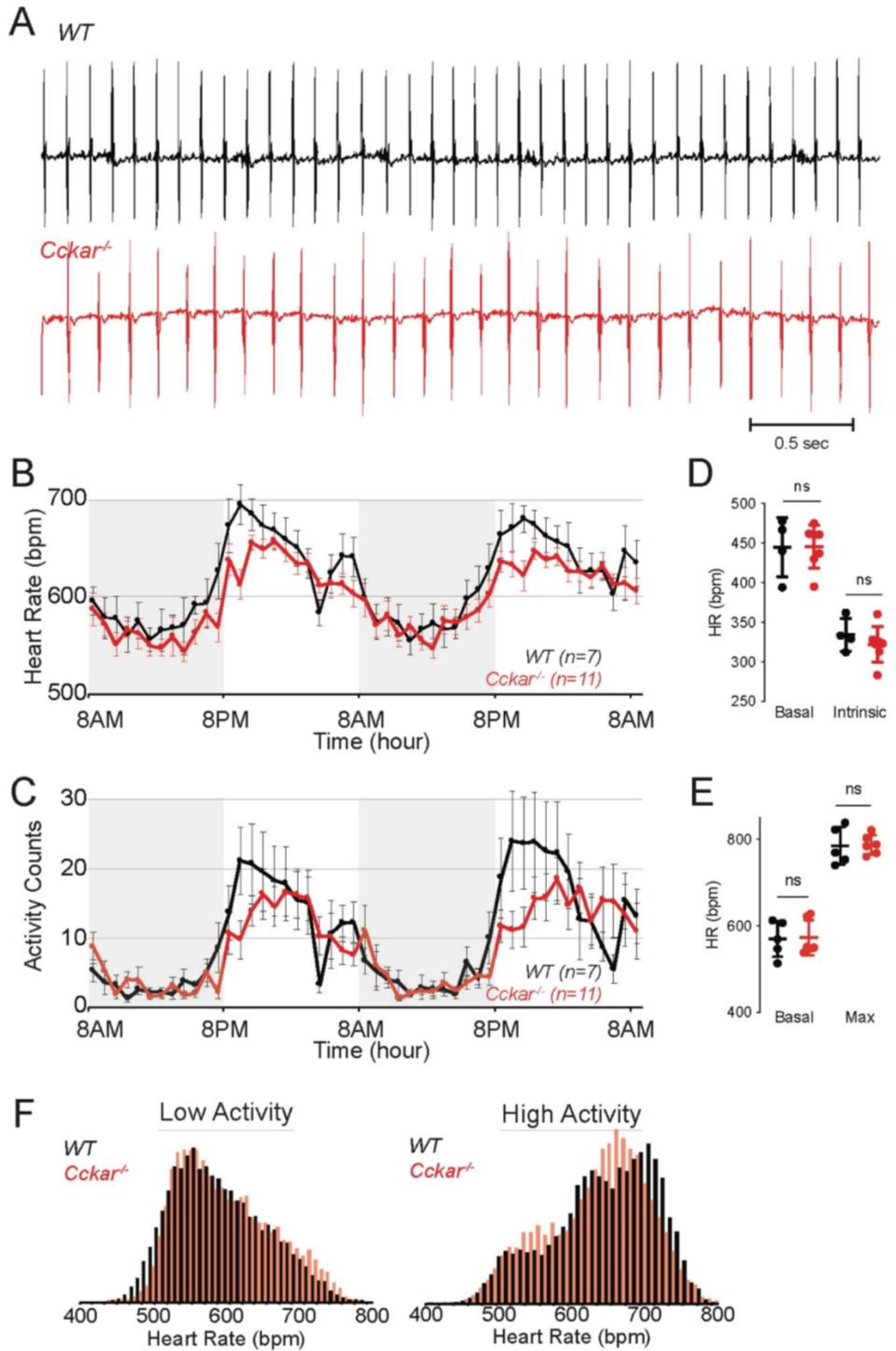
Heart Rhythm in *Cckar*^*-/-*^ Mice. (A) Electrocardiographic (ECG) tracings from awake unrestrained WT (top, black) and *Cckar*^*-/-*^ (bottom, red) adult mice. (B) Diurnal heart rate variation among 7 WT and 11 *Cckar*^*-/-*^adult mice implanted with ECG transmitters. Each point shows mean +/-SEM. (C) Activity counts for 7 WT and 11 *Cckar*^*-/-*^adult mice implanted with transmitters. Each point shows mean +/-SEM. (D) Heart rate before (baseline) and after atropine and propranolol injection (A/P) in 4 WT (black) and 7 *Cckar*^*-/-*^mice (red) under anesthesia. Error bars denote standard deviation. (E) Heart rate before (baseline) and after isoproterenol injection (Max) in 5 WT (black) and 6 *Cckar*^*-/-*^ mice (red). Error bars denote standard deviation. A Mann-Whitney test was used to test for significance with P < 0.05 deemed significant. ‘ns’ denotes non-significant. (F) Averaged histograms of heart rate during low activity period (left) and high activity period (right) in 7 WT (black) and 11 *Cckar*^*-/-*^mice (red) with 10 beat per minute bins.

### 3.5 sCCK-8 slows spontaneous firing in isolated adult PCs

To test whether sCCK-8 affects electrophysiological properties of PCs, we isolated PCs from adult WT C57Bl6 mice and measured action potentials in whole cell current clamp mode using the perforated patch technique. Individual spontaneously beating mouse pacemaker cells were exposed to different concentrations of sCCK-8 during AP recording. In the presence of 1nM isoproterenol, commonly used in isolated PCs to approximate behavior under physiological sympathetic tone, perfusion of sCCK-8 slowed firing rate in some but not all PCs (Figure 4A, B). The onset of the effect varied but generally occurred within one minute of exposure and was largely complete by 3 minutes. Firing rate did not recover to baseline after washing out the sCCK-8. None of the PCs isolated from adult *Cckar*^*-/-*^ mice exposed to 5 nM and 10 nM sCCK-8 exhibited an effect on firing rate, demonstrating that the negative chronotropic effect of sCCK-8 requires the presence of CCK_A_R (Figure 4C, D). The effect size varied, with statistically significant 10-20% slowing of firing rate at dosages of 2.5 nM (*n*=7), 10nM (*n*=6), and 20 nM (*n*=4) of s-CCK-8, and a non-significant effect observed at 1nM (*n*=5) and 5nM (*n*=6). (Figure 4E, F). Notably, a subset of spontaneously firing PCs did not respond to s-CCK8, as reflected in the broad distribution of percentage change in firing rate, possibly due to heterogeneity of CCK_A_R expression within the population of pacemaker cells.

**Figure 4.**
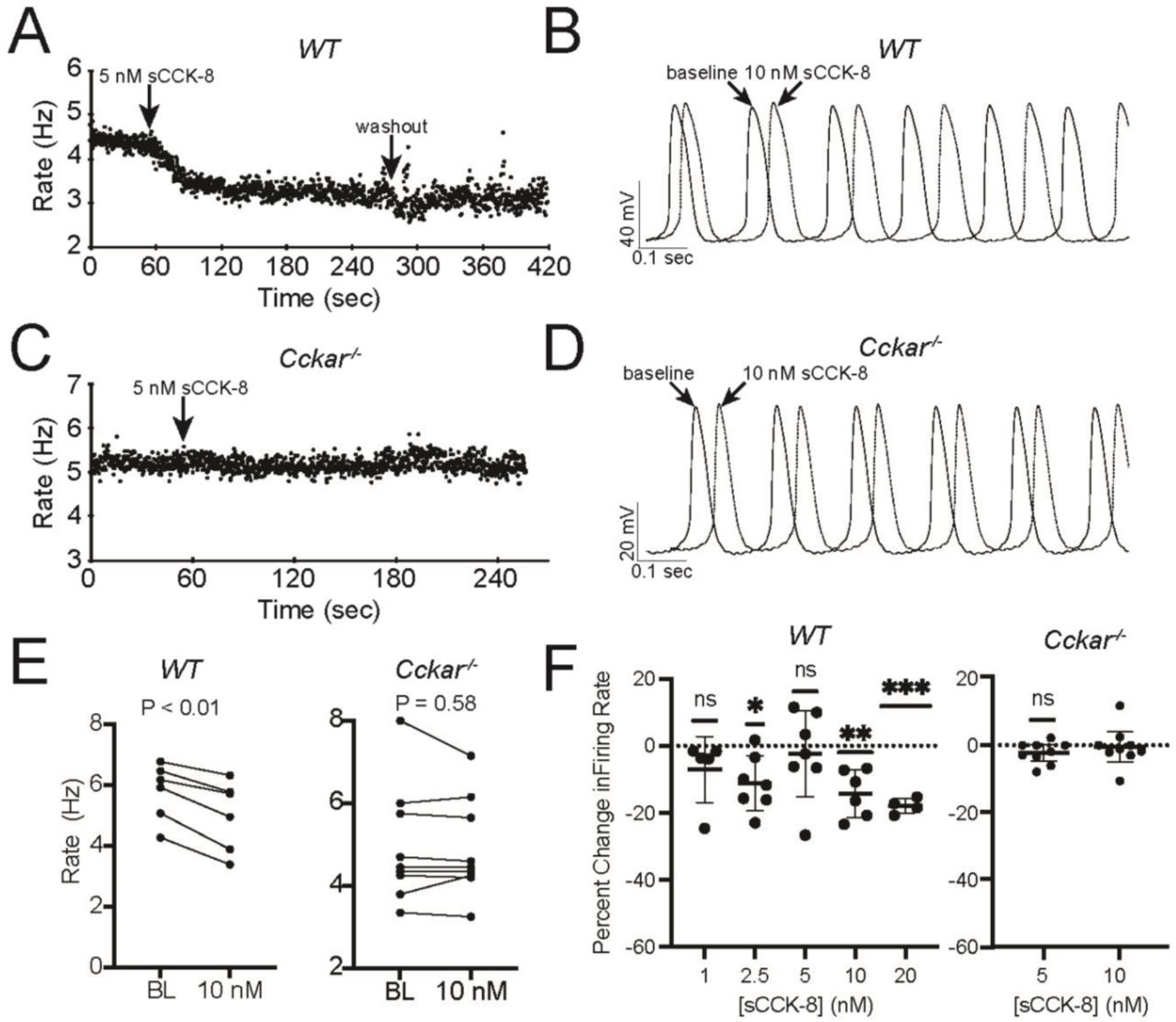
sCCK-8 Reduces Spontaneous Firing Rate of Pacemaker Cells. (A) Spontaneous firing rate for an isolated adult WT pacemaker cell before and during perfusion of 5 mM sCCK-8, and after washout. (B) Superimposed spontaneous action potentials from an adult WT pacemaker cell firing spontaneously at baseline (solid line) and after perfusion with 10 nM sCCK-8 (dashed line). (C) Spontaneous firing rate for an isolated adult *Cckar*^*-/-*^ pacemaker cell before and during perfusion with 5 nM sCCK-8. (D) Superimposed spontaneous action potentials from an adult *Cckar*^*-/-*^ pacemaker cell at baseline (solid line) and during perfusion with 10 mM sCCK-8 (dashed line). (D) Change in spontaneous firing rate of (E) adult WT (n = 6) and (F) adult *Cckar*^*-/-*^ (n = 9) pacemaker cells at baseline (BL) and 1 minute after perfusion of 10 nM sCCK-8. Statistical comparison was made with a paired T-test with the indicated P values. (F) Percent change in spontaneous firing rate for individual adult WT pacemaker cells 1 minute after perfusion of 1 nM (n=5), 2.5 nM (n=7), 5 nM (n=7), 10 nM (n=6), or 20 nM (n=4) sCCK-8. Statistical comparison between each group and 0% change (null hypothesis) was made with a one sample t-test (‘ns’ denotes not significant, ‘*” denotes P < 0.05, ‘**’ denotes P < 0.01, and ‘***’ denotes P < 0.001) (G) Percent change in firing rate in single adult *Cckar*^*-/-*^ pacemaker cells before and after perfusion with 5 nM (n=9) or 10 nM (n=9) sCCK-8. Statistical comparison between each group and 0% change (null hypothesis) was made with a one sample t-test Error bars indicate standard deviation throughout.

Considering the small effect size and high variability from cell-to-cell, we hypothesized that isoproterenol might have blunted the negative chronotropic response to sCCK-8, so we repeated these experiments using 5 nM sCCK-8 in the absence of isoproterenol. As expected, we observed a larger and consistent negative chronotropic effect of 5 nM sCCK-8 on firing rate without isoproterenol present, suggesting that sCCK-8 and isoproterenol activate pathways with opposing effects on the determinants of firing rate (Figure 5).

**Figure 5.**
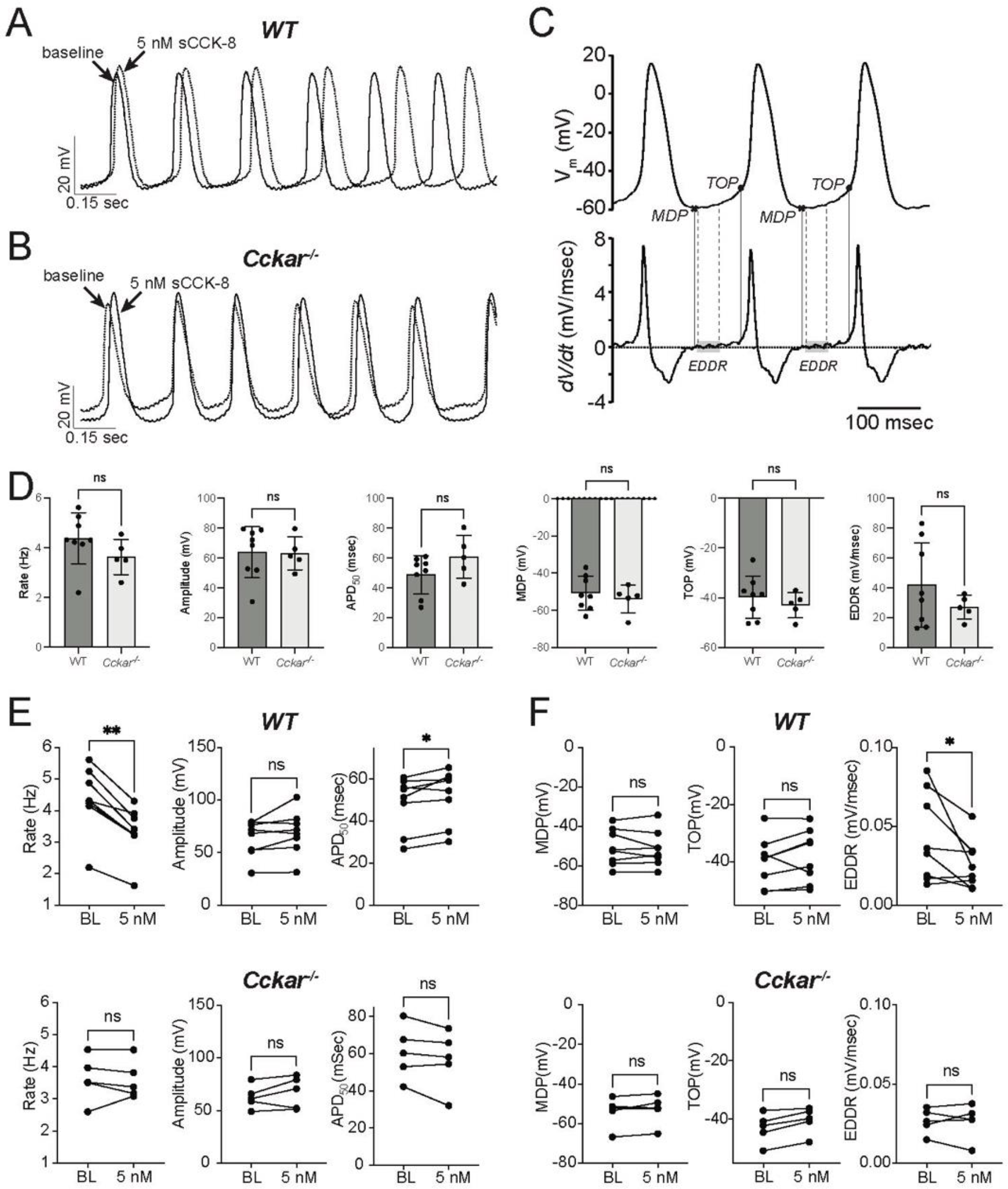
sCCK-8 Reduces Early Spontaneous Depolarization in Pacemaker Cardiomyocytes. (A) Slowing of spontaneous firing in response to sCCK-8 perfusion in a WT pacemaker cardiomyocyte (PC) in the absence of isoproterenol (B) Lack of response of a PC from a Cckar-/-SAN (C). Definition of action potential parameters. The top tracing, *V*_*m*_*(t)*, shows spontaneous action potentials (APs) from an isolated adult pacemaker cell and the bottom tracing shows the first derivative (*dV/dt*). The minimum diastolic potential (MDP) was defined as the minimum potential recorded between action potentials; Take off potential (TOP) was defined as the membrane potential at which dV/dt reach 10% of its maximum value; and early diastolic depolarization rate (EDDR) was defined as the average value of *dV/dt* over the shaded area in the bottom panel (from 10% to 50% of the MDP-to-TOP interval). (D) AP parameters including Firing Rate, Amplitude, APD_50_, MDP, TOP, and EDDR were measured under baseline conditions in WT and *Cckar*^*-/-*^ PCs (*n* = 8 and *n* = 5, respectively), and 3 minutes after perfusion with 5 nM sCCK-8. Statistical comparisons between baseline values and between pre- and post-sCCK-8 values were made with a Mann-Whitney test (‘ns’ denotes non-significant, ‘*’ denotes P < 0.05, ‘**’ denotes P < 0.01).

To better define the electrophysiological mechanism for the reduction in firing rate, AP morphology was quantified in the presence of sCCK-8 in the presence and absence of isoproterenol. Action potential duration at 50% repolarization (APD50, msec), actional potential amplitude (mV), maximum diastolic potential (MDP, mV), takeoff potential (TOP, mV), and early diastolic depolarization (EDDR, mV/msec) were determined at baseline and after perfusion with 5 or 10 nM sCCK-8. Of these parameters, we observed reductions in EDDR and an increase in APD50 in the presence of sCCK-8, suggesting a signaling mechanism that reduced spontaneous depolarization and affected repolarization without significantly affecting other components of the pacemaker cell action potential (Figure 5). The decrease in EDDR also occurred in the presence of isoproterenol, although the effect was less pronounced (Supplementary Figure 5). Importantly, we did not observe changes in AP morphology in PCs from *Cckar*^*-/-*^ mice exposed to sCCK-8, again underscoring the dependence of the pharmacological effect on the expression of CCK_A_R. Notably, there were no significant differences in firing rate or AP morphology between WT and KO PCs under baseline conditions.

## 4 DISCUSSION

Our main findings are (1) mammalian PCs express functional CCK_A_R, (2) CINS neurons and SAN fibroblasts express CCK, and (3) sCCK-8 slows PC firing rate by reducing the slope of early diastolic depolarization. Taken together, these data provide evidence for a novel signaling system in PCs that could affect mammalian SAN function through direct effects on automaticity.

### 4.1 GPCR Expression in the SAN

Analysis of existing RNA sequencing datasets identified several GPCRs that are enriched in SAN tissue and expressed in both SAN and atrial tissue, including orphan GPCRs and GPCRs with defined ligands but no known functions in regulating heart rhythm. Ligands for several of these GPCRs, including *Sstr4, S1pr3, Gcgr*, and *Adcyap1r1* trigger defined chronotropic responses but the molecular mechanisms driving their specific actions on isolated pacemaker cells have been characterized only for some^23^. Our findings of high-level and differential expression of these receptors in PCs suggest that chronotropic responses to these ligands are mediated by direct effects on PC automaticity. Further work will be required to define the specific electrophysiological effects of these signaling pathways on PCs in physiological contexts and to identify the relevant sources of their ligands. Importantly, leading computational models of the PC action potential have yet to incorporate dynamic signaling through simultaneously activated receptors for these various ligands, and the potential roles of these signaling pathways in long-term versus short-term SAN physiology have not been determined. The catalog of GPCRs presented here could form a foundation for such efforts and for deeper exploration of interaction between the CINS and SAN beyond the canonical signaling pathways involved in autonomic regulation of the heartbeat.

### 4.2 Expression of Cholecystokinin Signaling Pathway Components in the SAN

Recent studies have explored the functional neuroanatomy of the cardiac intrinsic nervous system with particular attention to innervation of the SAN. In a recent single cell qPCR analysis of porcine heart, CCK-expressing neurons were identified as a subset of right atrial ganglionated plexus neurons and included cells that projected directly to the SAN ^24^. Using a *Cck*^*Cre*^ mouse line to drive reporter gene expression, we uncovered similar findings in mice, with Cre-expressing neurons observed in the RAGP and in projections within the SAN, findings that we confirmed with analysis of scRNA-seq data. An additional finding of note is the presence of CCK-expressing fibroblasts within the SAN which has not been previously described and raises the possibility of paracrine signaling from SAN fibroblasts to pacemaker cells. The stimulus for secretion and whether such signaling might play a role in sinus node dysfunction remain to be determined. However, the deep conservation of expression of the CCK signaling system among mammals suggests a conserved functional role in SAN function and in autonomic regulation of heart rhythm.

### 4.3 Role of Cholecystokinin Signaling in Regulating Heart Rhythm

Although *Cckar*^*-/-*^ mice had modestly slower heart rates than WT mice during high activity periods, this might have resulted from lower activity levels in these mice owing to the involvement of CCK_A_R in many central and peripheral neuronal and organ-specific signaling pathways that affect mouse behavior. Thus, reduced activity in *Cckar*^*-/-*^ mice could have led to the slower heart rates, although we cannot exclude the opposite possibility that compromised SAN function reduced activity levels. However, our finding that *Cckar*^*-/-*^ mice do not exhibit sinus node dysfunction or changes in intrinsic heart rate after autonomic blockade suggests that CCK_A_R signaling is not required for SAN function under baseline conditions in adult mice. Related to this point, the magnitude of the sCCK-8 effect on isolated pacemaker cells is modest as assessed on a short time scale and could be counteracted by co-administration of isoproterenol, suggesting that the primary role of this pathway in autonomic regulation of SAN function may not be direct beat-to-beat regulation of heart rate. Alternatively, there may be redundant mechanisms that mitigate the loss of CCK_A_R in *Cckar*^*-/-*^mice.

### 4.4 Acute effects of Cckar Signaling on Isolated Pacemaker Cells

Several studies in rodent models have demonstrated that injection of sCCK-8 into an anesthetized animal or bolus injection of sCCK-8 to the heart *ex-vivo* can produce changes in heart rate. Our work confirms these results and extends the earlier findings by demonstrating that sCCK-8 acutely and directly reduces the firing rate of PCs by lowering the EDDR and lengthening the APD. While CCK_A_R is generally considered to signal preferentially via Gq ^25^, signaling via Gs ^26^, Gi ^27^, or G12/13 ^28, 29^ has been reported in different cellular contexts. Of note, activation of Gq specifically in murine SAN PCs using a genetically encoded optogenetic tool *increased* heart rate by ∼5% ^30^, while activation of Gq using DREADD technology resulted in a non-significant increase in heart rate ^31^, suggesting that the acute decrease in firing rate we observed may not result purely from signaling via Gq. Thus, the reduction in EDDR could have resulted from reduction in cyclic adenosine monophosphate (cAMP) levels as might be expected with signaling via Gi. The finding that the magnitude of the effect could be blunted by co-administration of isoproterenol, an activator of Gs, would support such a hypothesis.

### 4.5 Limitations

Our study does not directly address the possibility that CCK_A_R signaling affects SAN properties not related to acute physiological response. In addition, we used a global loss of function model, which confounds our ability to define the physiological consequences of perturbing CCK_A_R signaling specifically in PCs. Despite these limitations, the present study demonstrates functional expression of CCK_A_R in PCs and establishes a foundation for future work on this signaling pathway and its relevance to heart rhythm regulation.

## Acknowledgements

The authors are grateful to Huiliang Qui for assistance with preparing materials for physiology experiments, and to Gagandeep Chouhan for assistance with genotyping and mouse colony management.

