## Supplementary Materials for "Cholecystokinin-A Signaling Regulates Automaticity of Pacemaker Cardiomyocytes"

### Cholecystokinin-A Signaling Regulates Pacemaker Cardiomyocyte Automaticity and Shortens Sinus Node Recovery Time

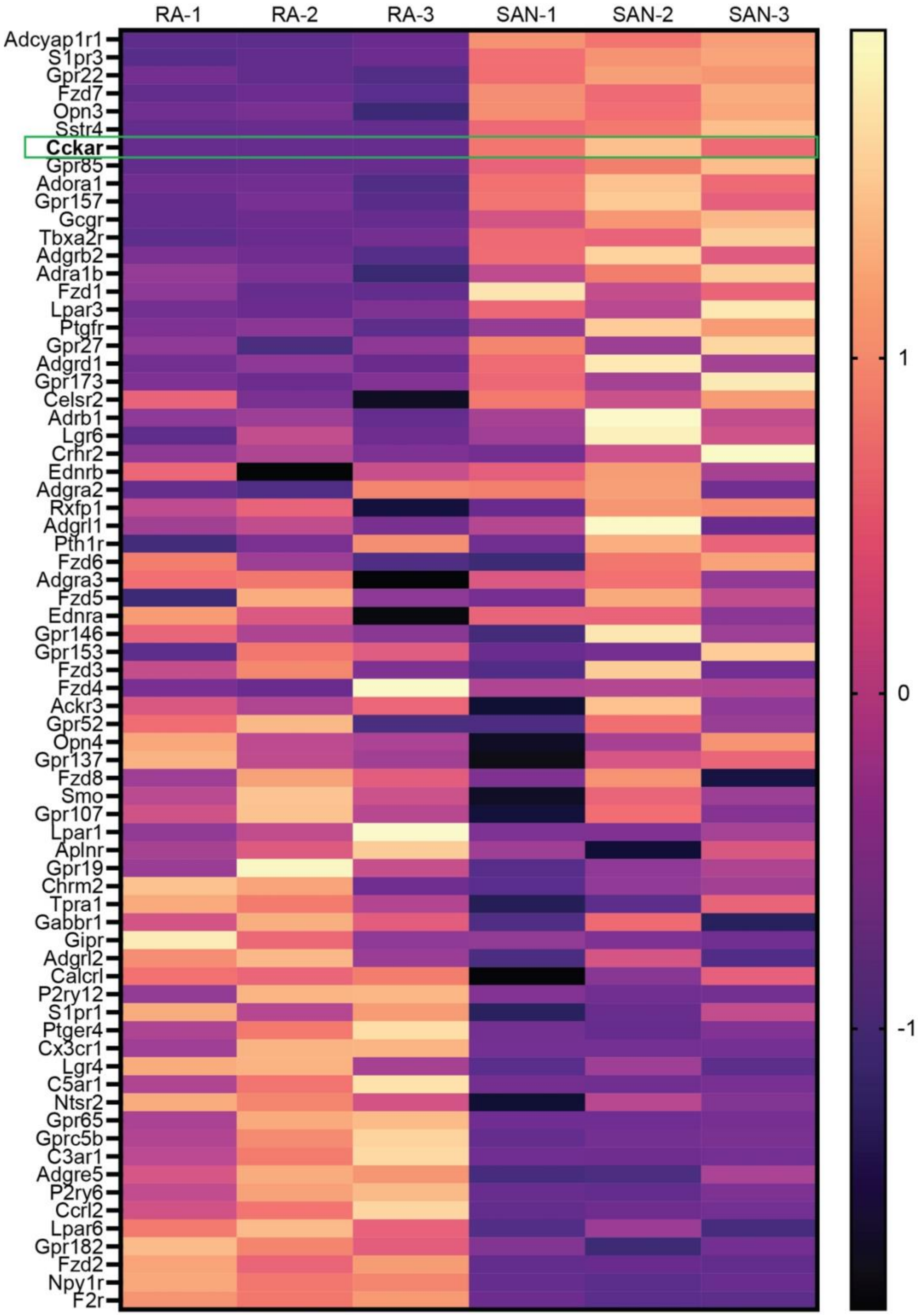


**Supplementary Figure 1. Differential Expression of GPCRs in Mammalian Sinoatrial Node.**

Heatmap demonstrates differentially expressed GPCRs from a previously published dataset [1] with 3 biological replicates of right atrial (RA) cardiomyocytes and sinoatrial node (SAN) pacemaker cardiomyocytes. Only receptors with significant differential expression (FDR < 0.05) are shown. *Cckar* is highlighted.

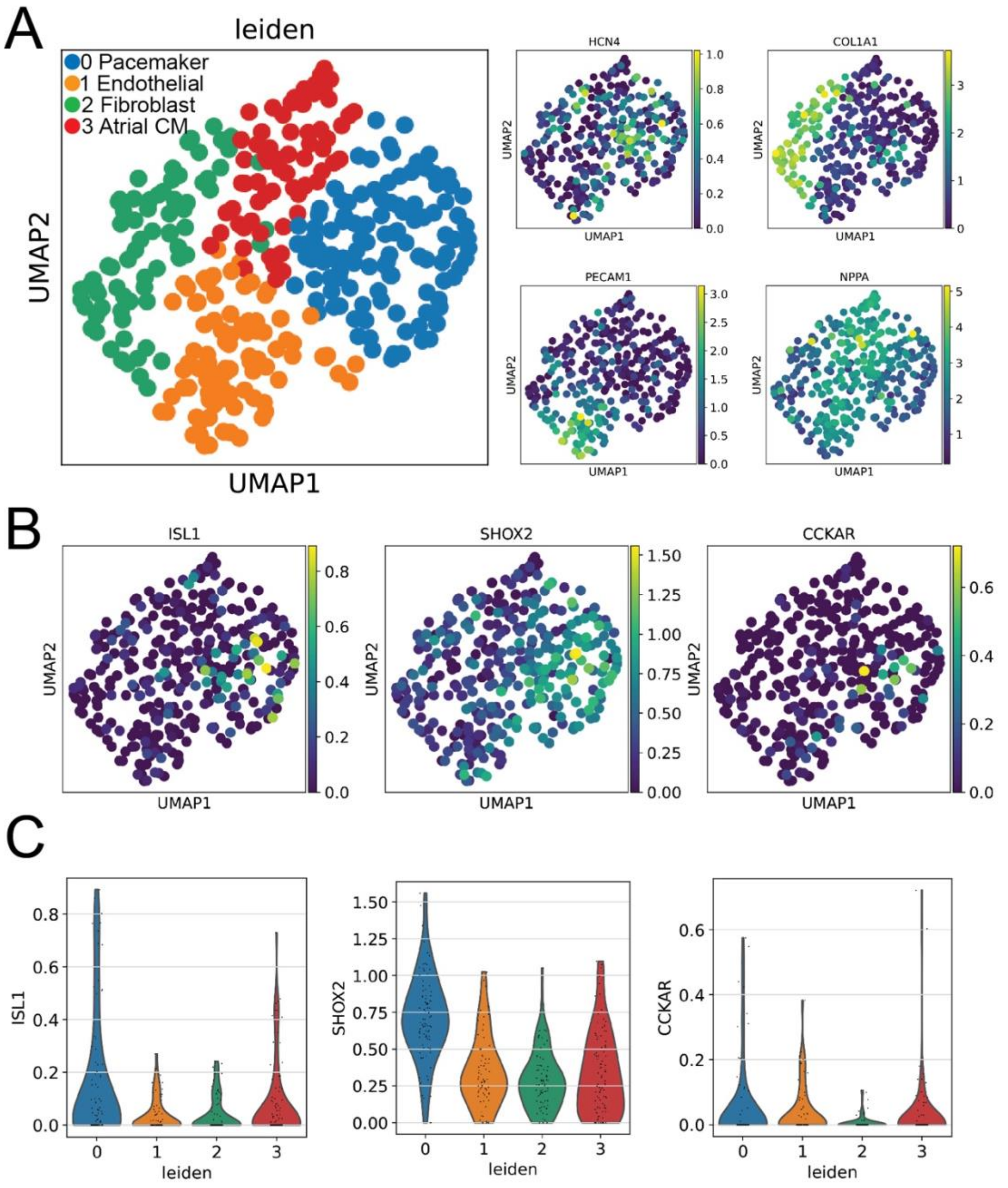


**<u>Supplementary Figure 2. Expression of Cckar in Rabbit Sinoatrial Node</u>**. (A). Uniform Manifold Approximation Projection (UMAP) plot of leiden clustering analysis of single cell RNA sequencing data derived from rabbit sinoatrial node tissue[2] demonstrated an *Hcn4*⁺ pacemaker cardiomyocyte (PC) population (cluster 0), as well as *Pecam*⁺ endothelial cells (cluster 1), *Col1a1*⁺ fibroblasts (cluster 2), and *Nppa*⁺ atrial cardiomyocytes (cluster 3) (B) UMAP plots demonstrating co-expression of established PC enriched genes *Shox2*, and *Isl1* alongside *Cckar* in cluster 0. (C) Quantification of expression of pacemaker genes Isl1, Shox2, and Cckar in different clusters.

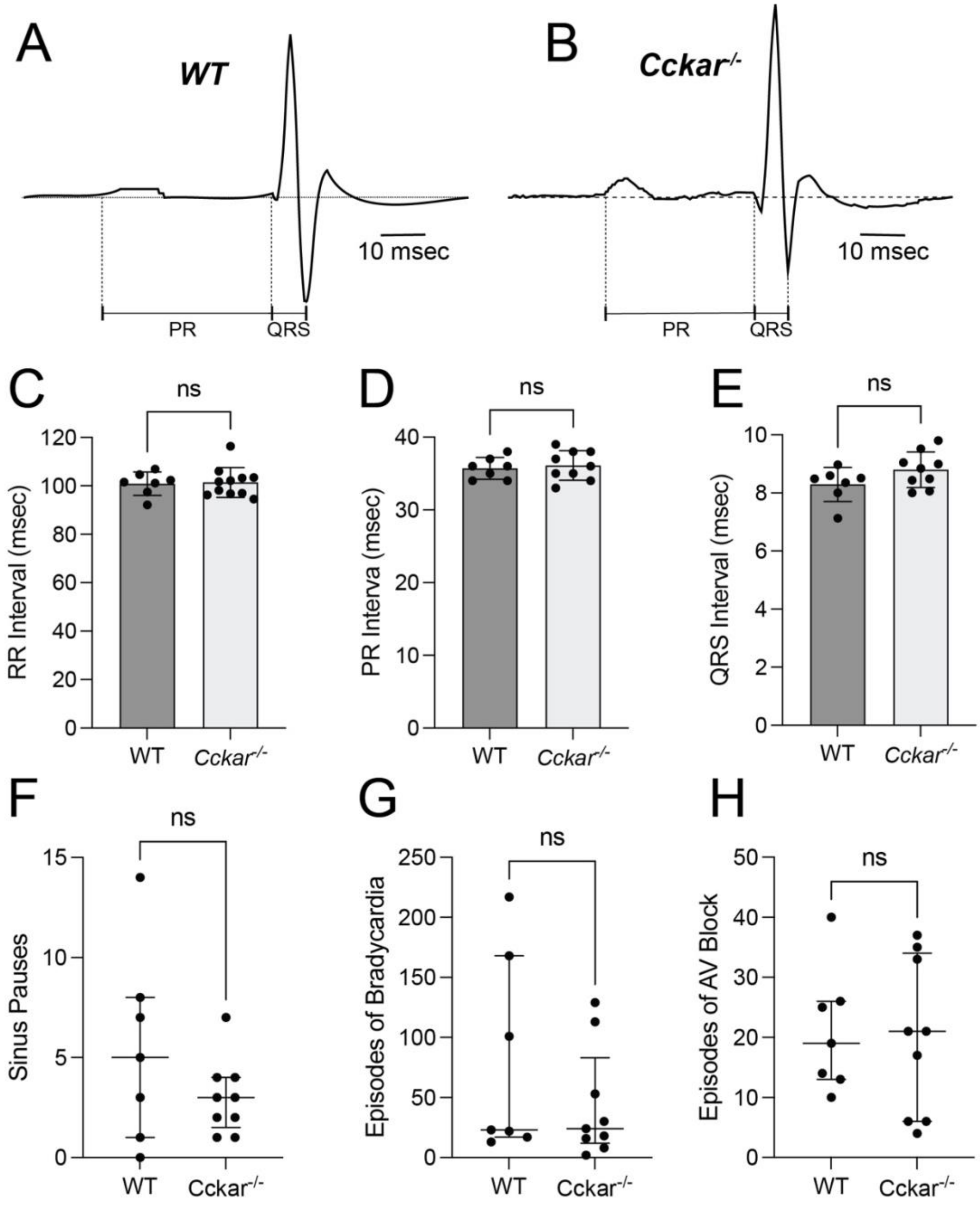


**<u>Supplementary Figure 3. Electrocardiographic Intervals and Arrhythmias in WT and *Cckar*$^{-/-}$ Mice</u>**. Averaged electrocardiogram (ECG) tracings from representative (A) WT and (B) *Cckar*$^{-/-}$ mice with ECG intervals indicated. (C) Quantification of RR, (D) PR, and (E) QRS intervals from WT (*n* = 7) and *Cckar*$^{-/-}$ (*n* = 9) mice. Sinus pauses (F), episodes of sinus bradycardia (G), and episodes of AV block (H) were quantified and summed for each mouse over 48 hours of continuous recording for WT (*n* = 7) and *Cckar*$^{-/-}$ (*n* = 9) mice. For electrocardiographic intervals, the line indicates the mean and errors bars indicate standard deviation; for arrhythmia counts, the line indicates the median and the error bars indicate interquartile intervals. For both intervals and arrhythmia episodes, Mann-Whitney tests were used to assess significance (‘ns’ denotes not significant, ‘*’ denotes $P < 0.5$).

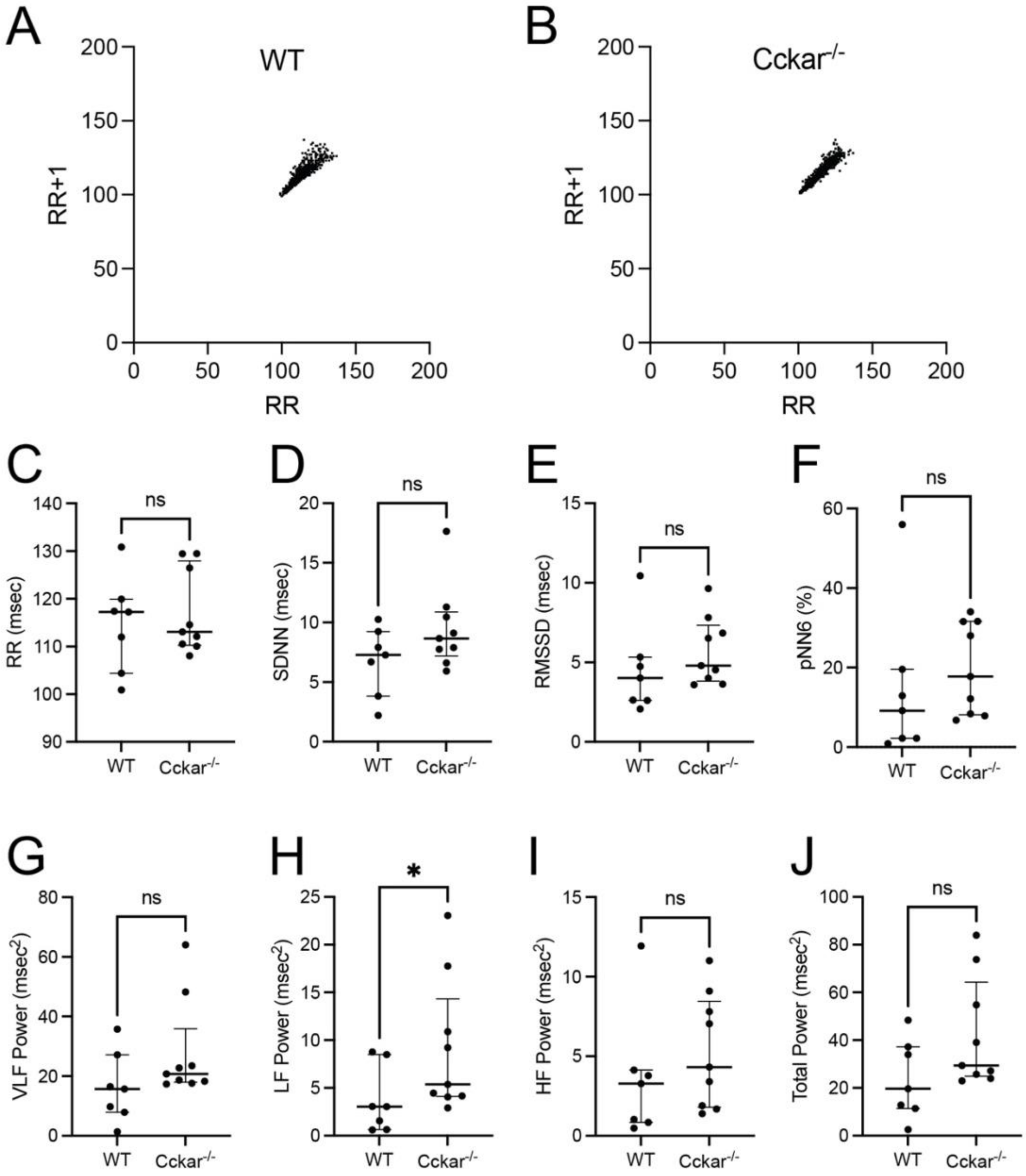


**Supplementary Figure 4. Heart Rate Variability in WT and *Cckar*$^{-/-}$ Mice**. Poincare plots are shown for a representative WT (A) and (B) *Cckar*$^{-/-}$ littermate during low activity conditions. The following parameters were determined from EKG tracings acquired during normal rhythm from implanted telemetry devices in freely moving unanesthetized WT ($n$ = 7) and *Cckar*$^{-/-}$ ($n$ = 9) littermates: (C) RR interval (D) standard deviation of the RR interval (SDNN), (E) Root mean square of successive differences between heartbeats (RMSSD), and (F) the percentage of successive RR intervals that differ by more than 6 msec (pNN6). Frequency domain analysis was used to calculate the contributions of (G) very low frequency (VLF), (H) low frequency (LF), (I) high frequency (HF) components to (J) the total power. The Mann-Whitney test was used for statistical comparison for all parameters and ‘ns’ denotes not significant.

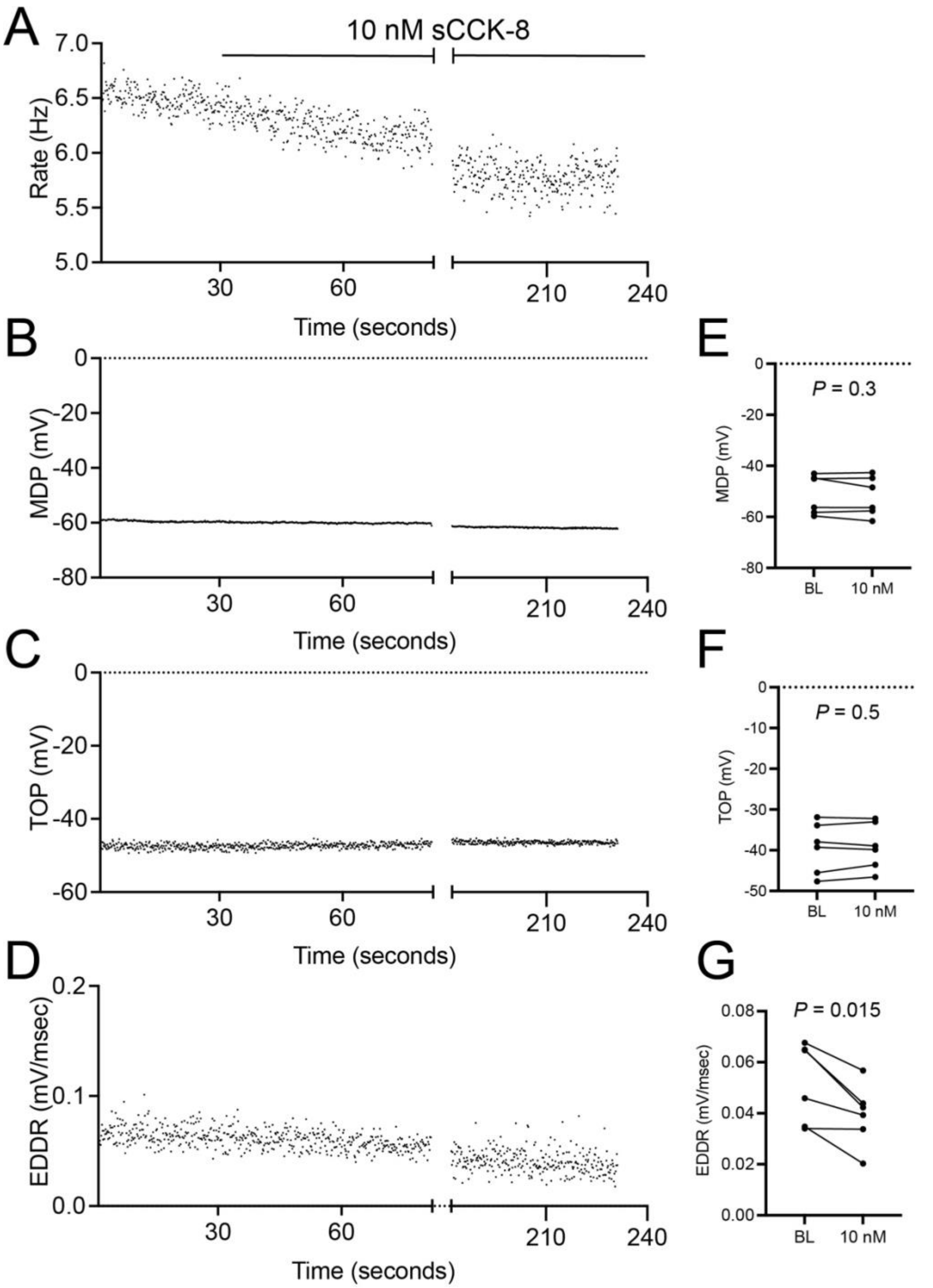


**Supplementary Figure 5. Effect of sCCK-8 on Pacemaker Cardiomyocyte Action Potential in the Presence of Isoproterenol.** (A). Beat-to-beat change in rate of a single pacemaker cardiomyocyte exposed to 10 nM sCCK-8 in the presence of 1 nM isoproterenol. Shown below are beat-to-beat changes in (B) minimum diastolic potential (MDP), (C) takeoff potential (TOP), and (D) early diastolic depolarization rate (EDDR) rate for the same cell. Shown on the right are before and after comparisons of (E) MDP, (F) TOP, and (G) EDDR for 5 different PCs. P value was calculated with a paired Mann-Whitney test.

#### Supplementary Table. Differential Expression Analysis for GPCRs: Pacemaker Cardiomyocytes (PC) vs Right Atrial Cardiomyocytes (RA)

| ensembl_gene_id | Gene | mRNA Expression (Transcripts per Million) | | | | | | zScore | | | | | | Avg z (PC) | P value | Rank | FDR |
|---|---|---|---|---|---|---|---|---|---|---|---|---|---|---|---|---|---|
| | | RA 1 | RA 2 | RA 3 | PC 1 | PC 2 | PC 3 | RA 1 | RA 2 | RA 3 | PC 1 | PC 2 | PC 3 | | | | |
| ENSMUSG00000029778 | Adcyap1r1 | 0.61 | 0.48 | 1.36 | 15.67 | 13.50 | 16.49 | -0.93 | -0.95 | -0.84 | 0.96 | 0.69 | 1.07 | 0.91 | 1.03E-04 | 2 | 0.0036 |
| ENSMUSG00000067586 | S1pr3 | 5.40 | 8.26 | 11.97 | 68.39 | 81.43 | 87.26 | -0.98 | -0.91 | -0.81 | 0.63 | 0.96 | 1.11 | 0.90 | 2.80E-04 | 4 | 0.0049 |
| ENSMUSG00000044067 | Gpr22 | 6.04 | 4.37 | 3.21 | 20.10 | 24.50 | 23.57 | -0.75 | -0.92 | -1.03 | 0.64 | 1.08 | 0.98 | 0.90 | 3.17E-04 | 5 | 0.0044 |
| ENSMUSG00000041075 | Fzd7 | 9.43 | 11.04 | 8.48 | 40.12 | 33.81 | 44.50 | -0.91 | -0.81 | -0.96 | 0.93 | 0.55 | 1.19 | 0.89 | 7.31E-04 | 6 | 0.0083 |
| ENSMUSG00000026525 | Opn3 | 3.82 | 4.08 | 2.18 | 10.63 | 9.39 | 11.50 | -0.77 | -0.71 | -1.18 | 0.92 | 0.61 | 1.13 | 0.89 | 1.12E-03 | 8 | 0.0094 |
| ENSMUSG00000037014 | Sstr4 | 0.11 | 0.39 | 0.00 | 17.54 | 19.93 | 27.52 | -0.88 | -0.86 | -0.89 | 0.54 | 0.74 | 1.36 | 0.88 | 2.03E-03 | 9 | 0.0149 |
| ENSMUSG00000029193 | Cckar | 0.00 | 0.00 | 0.05 | 26.49 | 37.85 | 24.04 | -0.88 | -0.88 | -0.87 | 0.70 | 1.38 | 0.55 | 0.88 | 2.29E-03 | 10 | 0.0149 |
| ENSMUSG00000048216 | Gpr85 | 0.15 | 0.24 | 0.23 | 2.85 | 3.50 | 4.59 | -0.90 | -0.86 | -0.86 | 0.47 | 0.80 | 1.35 | 0.88 | 2.49E-03 | 11 | 0.0145 |
| ENSMUSG00000042429 | Adora1 | 4.38 | 4.43 | 2.26 | 15.32 | 20.98 | 14.50 | -0.78 | -0.77 | -1.06 | 0.66 | 1.40 | 0.55 | 0.87 | 3.59E-03 | 12 | 0.0189 |
| ENSMUSG00000047875 | Gpr157 | 1.84 | 1.99 | 1.74 | 3.28 | 3.99 | 3.03 | -0.87 | -0.71 | -0.98 | 0.69 | 1.46 | 0.42 | 0.85 | 5.97E-03 | 13 | 0.0285 |
| ENSMUSG00000025127 | Gcgr | 0.95 | 3.30 | 1.41 | 35.58 | 59.08 | 68.85 | -0.88 | -0.81 | -0.87 | 0.24 | 1.00 | 1.32 | 0.85 | 6.02E-03 | 14 | 0.0262 |
| ENSMUSG00000034881 | Tbxa2r | 0.32 | 0.79 | 1.25 | 7.29 | 6.89 | 11.73 | -0.95 | -0.85 | -0.75 | 0.56 | 0.47 | 1.52 | 0.85 | 7.57E-03 | 16 | 0.0284 |
| ENSMUSG00000028782 | Adgrb2 | 2.83 | 2.49 | 1.58 | 7.67 | 11.42 | 6.90 | -0.69 | -0.78 | -1.02 | 0.57 | 1.55 | 0.37 | 0.83 | 1.16E-02 | 18 | 0.0382 |
| ENSMUSG00000050541 | Adra1b | 20.68 | 19.38 | 16.01 | 23.17 | 27.79 | 32.09 | -0.43 | -0.65 | -1.22 | 0.00 | 0.78 | 1.52 | 0.77 | 3.72E-02 | 24 | 0.0899 |
| ENSMUSG00000044674 | Fzd1 | 23.63 | 20.09 | 19.83 | 44.36 | 28.77 | 32.88 | -0.49 | -0.87 | -0.90 | 1.72 | 0.05 | 0.49 | 0.75 | 4.26E-02 | 26 | 0.0934 |
| ENSMUSG00000036832 | Lpar3 | 2.21 | 1.72 | 2.56 | 7.26 | 4.84 | 12.10 | -0.73 | -0.85 | -0.64 | 0.54 | -0.07 | 1.75 | 0.74 | 5.13E-02 | 31 | 0.0928 |
| ENSMUSG00000028036 | Ptgfr | 2.04 | 2.37 | 1.27 | 2.64 | 7.57 | 6.43 | -0.64 | -0.52 | -0.94 | -0.41 | 1.48 | 1.04 | 0.70 | 7.47E-02 | 35 | 0.1174 |
| ENSMUSG00000072875 | Gpr27 | 3.12 | 1.78 | 3.06 | 6.05 | 3.42 | 7.63 | -0.48 | -1.09 | -0.51 | 0.85 | -0.34 | 1.57 | 0.69 | 7.97E-02 | 36 | 0.1196 |
| ENSMUSG00000044017 | Adgrd1 | 1.01 | 1.96 | 0.49 | 6.38 | 11.28 | 2.91 | -0.73 | -0.50 | -0.85 | 0.58 | 1.76 | -0.26 | 0.69 | 8.16E-02 | 37 | 0.1169 |
| ENSMUSG00000056679 | Gpr173 | 2.40 | 2.12 | 2.45 | 4.41 | 3.05 | 6.53 | -0.64 | -0.81 | -0.62 | 0.54 | -0.26 | 1.79 | 0.69 | 8.28E-02 | 38 | 0.1133 |
| ENSMUSG00000068740 | Celsr2 | 3.42 | 2.21 | 1.24 | 3.69 | 3.04 | 3.99 | 0.47 | -0.69 | -1.64 | 0.74 | 0.10 | 1.03 | 0.62 | 1.37E-01 | 40 | 0.1741 |
| ENSMUSG00000035283 | Adrb1 | 15.90 | 17.71 | 10.81 | 19.09 | 47.15 | 22.44 | -0.49 | -0.35 | -0.89 | -0.24 | 1.95 | 0.02 | 0.58 | 1.80E-01 | 45 | 0.1999 |
| ENSMUSG00000042793 | Lgr6 | 19.13 | 26.59 | 20.16 | 23.69 | 40.38 | 27.58 | -0.93 | 0.04 | -0.79 | -0.33 | 1.84 | 0.17 | 0.56 | 1.97E-01 | 47 | 0.2055 |
| ENSMUSG00000003476 | Crhr2 | 5.84 | 6.54 | 5.47 | 5.29 | 7.21 | 10.99 | -0.49 | -0.16 | -0.67 | -0.75 | 0.15 | 1.92 | 0.44 | 3.31E-01 | 53 | 0.2997 |
| ENSMUSG00000022122 | Ednrb | 10.09 | 7.39 | 9.59 | 9.96 | 10.71 | 9.23 | 0.52 | -1.84 | 0.08 | 0.41 | 1.06 | -0.24 | 0.41 | 3.71E-01 | 54 | 0.3229 |

| ensembl_gene_id | Gene | mRNA Expression (Transcripts per Million) | | | | | | zScore | | | | | | Avg z (PC) | P value | Rank | FDR |
|---|---|---|---|---|---|---|---|---|---|---|---|---|---|---|---|---|---|
| | | RA 1 | RA 2 | RA 3 | PC 1 | PC 2 | PC 3 | RA 1 | RA 2 | RA 3 | PC 1 | PC 2 | PC 3 | | | | |
| ENSMUSG00000031486 | Adgra2 | 7.70 | 7.38 | 10.98 | 10.91 | 11.43 | 7.92 | -0.89 | -1.05 | 0.84 | 0.80 | 1.08 | -0.77 | 0.37 | 4.28E-01 | 56 | 0.3517 |
| ENSMUSG00000034009 | Rxfp1 | 35.59 | 39.31 | 23.78 | 28.85 | 43.41 | 42.60 | 0.00 | 0.47 | -1.50 | -0.86 | 0.99 | 0.89 | 0.34 | 4.63E-01 | 57 | 0.3654 |
| ENSMUSG00000013033 | Adgrl1 | 21.05 | 22.93 | 18.46 | 22.25 | 34.67 | 17.55 | -0.29 | 0.02 | -0.71 | -0.09 | 1.92 | -0.85 | 0.32 | 4.89E-01 | 58 | 0.3708 |
| ENSMUSG00000032492 | Pth1r | 4.52 | 5.53 | 9.15 | 5.37 | 9.78 | 8.11 | -1.16 | -0.70 | 0.94 | -0.77 | 1.22 | 0.47 | 0.31 | 5.16E-01 | 60 | 0.3700 |
| ENSMUSG00000022297 | Fzd6 | 7.68 | 5.75 | 4.50 | 4.28 | 7.59 | 8.26 | 0.77 | -0.34 | -1.06 | -1.19 | 0.72 | 1.11 | 0.21 | 6.59E-01 | 63 | 0.4390 |
| ENSMUSG00000029090 | Adgra3 | 8.95 | 9.06 | 5.80 | 8.53 | 8.97 | 7.56 | 0.63 | 0.71 | -1.84 | 0.30 | 0.65 | -0.46 | 0.16 | 7.32E-01 | 65 | 0.4620 |
| ENSMUSG00000045005 | Fzd5 | 3.80 | 6.03 | 4.43 | 4.23 | 5.99 | 4.92 | -1.18 | 1.21 | -0.50 | -0.72 | 1.17 | 0.02 | 0.16 | 7.44E-01 | 66 | 0.4510 |
| ENSMUSG00000031616 | Ednra | 89.43 | 76.86 | 41.64 | 79.98 | 79.66 | 62.64 | 1.04 | 0.30 | -1.76 | 0.48 | 0.47 | -0.53 | 0.14 | 7.71E-01 | 68 | 0.4424 |
| ENSMUSG00000044197 | Gpr146 | 12.78 | 11.03 | 10.11 | 8.55 | 16.02 | 10.64 | 0.49 | -0.19 | -0.54 | -1.14 | 1.73 | -0.34 | 0.08 | 8.65E-01 | 70 | 0.4694 |
| ENSMUSG00000042804 | Gpr153 | 8.63 | 13.96 | 12.77 | 8.87 | 9.24 | 16.42 | -0.94 | 0.72 | 0.35 | -0.86 | -0.75 | 1.48 | -0.04 | 9.31E-01 | 71 | 0.4850 |
| ENSMUSG00000007989 | Fzd3 | 10.04 | 10.95 | 9.26 | 8.82 | 11.66 | 9.12 | 0.06 | 0.86 | -0.63 | -1.03 | 1.49 | -0.76 | -0.10 | 8.42E-01 | 69 | 0.4393 |
| ENSMUSG00000049791 | Fzd4 | 7.45 | 7.04 | 14.08 | 8.73 | 8.88 | 8.82 | -0.68 | -0.84 | 1.94 | -0.17 | -0.11 | -0.14 | -0.14 | 7.71E-01 | 67 | 0.4027 |
| ENSMUSG00000044337 | Ackr3 | 4.13 | 3.48 | 4.45 | 1.49 | 5.72 | 3.05 | 0.29 | -0.17 | 0.51 | -1.56 | 1.40 | -0.47 | -0.21 | 6.60E-01 | 64 | 0.3506 |
| ENSMUSG00000118401 | Gpr52 | 8.45 | 10.72 | 3.11 | 3.16 | 8.60 | 5.35 | 0.60 | 1.31 | -1.09 | -1.08 | 0.64 | -0.38 | -0.27 | 5.66E-01 | 62 | 0.3013 |
| ENSMUSG00000021799 | Opn4 | 10.71 | 8.57 | 8.25 | 5.64 | 8.17 | 10.39 | 1.14 | -0.03 | -0.20 | -1.63 | -0.25 | 0.97 | -0.30 | 5.20E-01 | 61 | 0.2727 |
| ENSMUSG00000024958 | Gpr137 | 4.72 | 4.12 | 3.98 | 3.32 | 4.24 | 4.35 | 1.27 | -0.01 | -0.30 | -1.72 | 0.26 | 0.50 | -0.32 | 4.93E-01 | 59 | 0.2589 |
| ENSMUSG00000036904 | Fzd8 | 3.90 | 5.32 | 4.62 | 3.60 | 5.20 | 2.76 | -0.33 | 1.09 | 0.39 | -0.64 | 0.97 | -1.48 | -0.38 | 4.09E-01 | 55 | 0.2231 |
| ENSMUSG00000001761 | Smo | 12.83 | 17.56 | 13.41 | 7.71 | 14.56 | 11.83 | -0.05 | 1.41 | 0.13 | -1.62 | 0.48 | -0.35 | -0.50 | 2.63E-01 | 52 | 0.1468 |
| ENSMUSG00000000194 | Gpr107 | 7.43 | 8.64 | 7.16 | 5.71 | 7.85 | 6.65 | 0.19 | 1.39 | -0.08 | -1.52 | 0.61 | -0.59 | -0.50 | 2.61E-01 | 51 | 0.1433 |
| ENSMUSG00000038668 | Lpar1 | 2.21 | 3.96 | 11.38 | 1.44 | 1.53 | 2.95 | -0.45 | 0.01 | 1.98 | -0.65 | -0.63 | -0.26 | -0.51 | 2.45E-01 | 50 | 0.1324 |
| ENSMUSG00000044338 | Aplnr | 2.19 | 3.11 | 5.01 | 2.04 | 0.06 | 3.04 | -0.24 | 0.33 | 1.50 | -0.33 | -1.55 | 0.29 | -0.53 | 2.27E-01 | 49 | 0.1204 |
| ENSMUSG00000032641 | Gpr19 | 2.66 | 4.41 | 3.01 | 2.18 | 2.58 | 2.83 | -0.37 | 1.90 | 0.09 | -0.99 | -0.48 | -0.15 | -0.54 | 2.15E-01 | 48 | 0.1118 |
| ENSMUSG00000045613 | Chrm2 | 87.19 | 84.50 | 65.91 | 64.21 | 69.21 | 70.78 | 1.39 | 1.11 | -0.79 | -0.96 | -0.45 | -0.29 | -0.57 | 1.86E-01 | 46 | 0.0970 |
| ENSMUSG00000002871 | Tpra1 | 8.24 | 7.72 | 6.68 | 5.16 | 5.67 | 7.40 | 1.18 | 0.76 | -0.11 | -1.37 | -0.95 | 0.49 | -0.61 | 1.46E-01 | 44 | 0.0763 |
| ENSMUSG00000024462 | Gabbr1 | 4.73 | 6.29 | 4.97 | 2.71 | 5.24 | 2.27 | 0.23 | 1.24 | 0.39 | -1.06 | 0.56 | -1.35 | -0.62 | 1.39E-01 | 43 | 0.0712 |
| ENSMUSG00000030406 | Gipr | 31.30 | 16.10 | 4.12 | 4.56 | 2.04 | 0.69 | 1.81 | 0.53 | -0.48 | -0.44 | -0.65 | -0.77 | -0.62 | 1.37E-01 | 42 | 0.0685 |
| ENSMUSG00000028184 | Adgrl2 | 19.91 | 21.42 | 15.25 | 12.65 | 17.49 | 12.86 | 0.91 | 1.32 | -0.37 | -1.08 | 0.25 | -1.03 | -0.62 | 1.37E-01 | 41 | 0.0668 |
| ENSMUSG00000059588 | Calcrl | 5.38 | 5.17 | 5.56 | 1.95 | 3.68 | 5.07 | 0.65 | 0.50 | 0.78 | -1.79 | -0.56 | 0.43 | -0.64 | 1.18E-01 | 39 | 0.0577 |

| ensembl_gene_id | Gene | mRNA Expression (Transcripts per Million) | | | | | | zScore | | | | | | Avg z (PC) | P value | Rank | FDR |
|---|---|---|---|---|---|---|---|---|---|---|---|---|---|---|---|---|---|
| | | RA 1 | RA 2 | RA 3 | PC 1 | PC 2 | PC 3 | RA 1 | RA 2 | RA 3 | PC 1 | PC 2 | PC 3 | | | | |
| ENSMUSG00000036353 | P2ry12 | 1.16 | 6.35 | 6.36 | 0.64 | 0.17 | 0.30 | -0.44 | 1.28 | 1.28 | -0.62 | -0.77 | -0.73 | -0.71 | 7.07E-02 | 34 | 0.0374 |
| ENSMUSG00000045092 | S1pr1 | 21.54 | 17.87 | 21.06 | 14.37 | 15.67 | 18.25 | 1.20 | -0.09 | 1.03 | -1.32 | -0.86 | 0.04 | -0.71 | 6.66E-02 | 33 | 0.0343 |
| ENSMUSG00000039942 | Ptger4 | 2.22 | 4.89 | 7.58 | 0.63 | 0.21 | 0.97 | -0.18 | 0.74 | 1.66 | -0.73 | -0.87 | -0.61 | -0.74 | 5.14E-02 | 32 | 0.0257 |
| ENSMUSG00000052336 | Cx3cr1 | 2.14 | 9.65 | 9.70 | 0.22 | 0.19 | 0.21 | -0.33 | 1.27 | 1.28 | -0.74 | -0.74 | -0.74 | -0.74 | 5.04E-02 | 30 | 0.0252 |
| ENSMUSG00000050199 | Lgr4 | 26.92 | 27.19 | 19.89 | 16.30 | 19.50 | 16.42 | 1.20 | 1.26 | -0.23 | -0.97 | -0.31 | -0.94 | -0.74 | 4.97E-02 | 29 | 0.0240 |
| ENSMUSG00000049130 | C5ar1 | 7.80 | 17.44 | 29.14 | 0.78 | 0.59 | 1.24 | -0.15 | 0.68 | 1.69 | -0.75 | -0.77 | -0.71 | -0.74 | 4.90E-02 | 28 | 0.0227 |
| ENSMUSG00000020591 | Ntsr2 | 5.03 | 4.61 | 3.89 | 1.83 | 3.55 | 2.95 | 1.20 | 0.84 | 0.21 | -1.56 | -0.08 | -0.60 | -0.75 | 4.54E-02 | 27 | 0.0202 |
| ENSMUSG00000021886 | Gpr65 | 1.28 | 4.44 | 4.83 | 0.00 | 0.00 | 0.17 | -0.22 | 1.17 | 1.35 | -0.79 | -0.79 | -0.72 | -0.77 | 3.72E-02 | 25 | 0.0164 |
| ENSMUSG00000008734 | Gprc5b | 4.40 | 6.63 | 8.12 | 2.77 | 3.11 | 3.17 | -0.14 | 0.88 | 1.56 | -0.88 | -0.72 | -0.70 | -0.77 | 3.65E-02 | 23 | 0.0159 |
| ENSMUSG00000040552 | C3ar1 | 8.81 | 17.87 | 27.40 | 0.72 | 0.60 | 1.36 | -0.06 | 0.76 | 1.62 | -0.79 | -0.80 | -0.73 | -0.77 | 3.32E-02 | 22 | 0.0136 |
| ENSMUSG00000002885 | Adgre5 | 18.87 | 27.26 | 25.36 | 6.62 | 6.84 | 14.75 | 0.25 | 1.20 | 0.99 | -1.13 | -1.10 | -0.21 | -0.81 | 1.74E-02 | 21 | 0.0066 |
| ENSMUSG00000048779 | P2ry6 | 3.23 | 6.89 | 7.70 | 0.22 | 0.04 | 0.89 | 0.02 | 1.09 | 1.33 | -0.86 | -0.92 | -0.67 | -0.82 | 1.65E-02 | 20 | 0.0058 |
| ENSMUSG00000043953 | Ccrl2 | 21.36 | 25.65 | 33.57 | 11.78 | 12.62 | 13.16 | 0.19 | 0.68 | 1.58 | -0.90 | -0.80 | -0.74 | -0.82 | 1.63E-02 | 19 | 0.0052 |
| ENSMUSG00000033446 | Lpar6 | 4.06 | 4.60 | 3.81 | 2.49 | 3.09 | 2.40 | 0.73 | 1.34 | 0.45 | -1.03 | -0.36 | -1.13 | -0.84 | 9.18E-03 | 17 | 0.0027 |
| ENSMUSG00000058396 | Gpr182 | 4.76 | 3.88 | 3.11 | 1.38 | 0.35 | 1.11 | 1.34 | 0.83 | 0.39 | -0.61 | -1.19 | -0.76 | -0.85 | 6.26E-03 | 15 | 0.0017 |
| ENSMUSG00000050288 | Fzd2 | 7.89 | 6.74 | 7.78 | 4.17 | 4.26 | 4.15 | 1.12 | 0.49 | 1.06 | -0.90 | -0.86 | -0.91 | -0.89 | 8.97E-04 | 7 | 0.0004 |
| ENSMUSG00000036437 | Npy1r | 28.31 | 23.18 | 24.82 | 3.55 | 2.68 | 3.91 | 1.14 | 0.72 | 0.85 | -0.89 | -0.96 | -0.86 | -0.90 | 1.44E-04 | 3 | 0.0001 |
| ENSMUSG00000048376 | F2r | 58.44 | 52.68 | 59.36 | 18.10 | 14.47 | 15.12 | 0.98 | 0.72 | 1.02 | -0.81 | -0.97 | -0.94 | -0.91 | 6.59E-05 | 1 | 0.0001 |

#### SUPPLEMENTARY METHODS

##### *Animal Handling and Maintenance*

Mouse studies were performed in accordance with IACUC-approved protocols at the University of California, San Francisco. Mouse lines were maintained on C57Bl6. *Cckar*$^{tm1Kpn}$ (hereafter, *Cckar*$^{-/-}$) mice were generated by Alan Kopin et al. and were obtained from Nirao Shah's lab [3]. *Cck*$^{tm1.1(cre)Zjh}$ (hereafter, *Cck*$^{Cre}$) mice were obtained from Jackson Labs. *Gt(ROSA)26Sor*$^{tm4(ACTB\text{-}tdTomato,\text{-}EGFP)Luo}$ (hereafter, *ROSA*$^{mT/mG}$ mice were generated by Muzumdar et al. and were obtained from Brian Black's lab [4].

##### *Analysis of Bulk RNA Sequencing Data*

To identify G-protein coupled receptors (GPCRs) that were expressed in pacemaker cardiomyocytes (PCs) and right atrial cardiomyocytes (RACMs), respectively, RNA-seq data from neonatal sorted mouse PCs and RACMs were downloaded, and the list of genes expressed in each tissue as defined by average transcripts per million across 3 biological replicates greater than 3 was intersected with an annotated database of from the International Union of Basic and Clinical Pharmacology. For each GPCR identified, a false discovery rate was calculated for differential expression was determined using the method of Benjamini-Hochberg.

##### *Single Cell RNA Sequencing Analysis*

Mouse E16.5 SAN s 10x Genomics Mouse single-cell RNA-sequencing files were generated by Goodyer et al. [5]and downloaded from GEO (GSE132658). Adult rabbit single-cell RNA-seq data were generated by Liang et al.[2] and were downloaded as fastq files from the NCBI Sequence Read Archive (PRJNA531288). Fastq files from the rabbit dataset were trimmed using Fastp[6] and aligned to the oryCun2 assembly (UCSC genome browser) using Hisat2[7]. Finally, an expression matrix was constructed for each cell using featureCounts[8]. Both mouse and rabbit datasets were filtered, clustered, and analyzed using Scanpy[9] to identify the cell types expressing both *Cckar* and *Cck*.

##### *Quantitative PCR*

The SAN and RA regions from 3 adult hearts were manually dissected using anatomical landmarks to delineate the inter-caval region. For each sample, total RNA from SAN and right atrium tissue was extracted using TRIzol reagent (Life technologies, CA, USA) and RNeasy micro kit (QIAGEN) according to the manufacturer's instructions. 0.2 micrograms of total RNA were used to synthesize the first-strand cDNA with Superscript III first-strand synthesis kit (Invitrogen). Real time qPCR was performed using Taqman fast advanced master mix (Thermo Fisher Scientific, USA). GAPDH was used as an endogenous control with the ΔΔCT method for comparison of SAN and right atrial tissue. A Mann-Whitney test was used for significance with $P < 0.05$ deemed significant.

##### *Immunohistochemistry*

Freshly isolated hearts were perfused with PBS, fixed in 4% PFA overnight at 4°C, washed in PBS, incubated in 15% sucrose followed by 30% sucrose, and then embedded in Optimal Temperature Cutting medium (Tissue Tek). 7.0 µm sections were cut with a Leica CM3050S cryostat. For antibody staining, sections were blocked in 0.1% saponin and 1% casein solution (Thermo Fisher, Catalog#37582) for 1 hour at room temperature and incubated with primary antibody overnight at 4°C (rabbit anti-Hcn4, Alomone Labs, APC-052, 1:300; chicken anti-GFP, Aves Labs GFP-1010, 1:300; goat anti-TdTomato, Origene, AB8181-200, 1:400). Sections were incubated with appropriate AlexaFluor conjugated secondary antibodies at 1:1000 for 1 hour at room temperature. DAPI was used to visualize the nuclei.

For acetylcholinesterase staining, sections were briefly washed with PBS, incubated in the reaction solution, which contains (in 10 ml of volume: acetyl thiocholine iodide 5mg; 0.1M sodium acetate (pH 6.0) 6.5ml; 0.1M sodium citrate 0.5ml; 30mM copper sulphate 1.0ml; 5mM potassium ferricyanide 1.0ml) for 2 hours at room temperature, rinsed with water and mounted.

##### *Transmitter Implantation and Analysis of Heart Rhythm Data*

Adult mice (8-12 weeks old) underwent subcutaneous implantation of telemetry devices (ETA-F10, Data Science International, MN) under anesthesia according to the manufacturer's instructions. After at least 7 days of recovery, continuous ECG signals were sampled using Ponemah software (DSI) at 1.0 kHz on freely moving and conscious mice for 48 hours per mouse. Maximum heart rate was ascertained after injection of 1.0 µg of isoproterenol. To determine the intrinsic heart rate, a separate cohort of mice were injected with 3.0 mg/kg atropine and 10.0 mg/kg propranolol intraperitoneally under anesthesia with 1% isoflurane. Anesthetized ECG signals were recorded with a Power Lab and Animal BioAmp (AD Instruments) and analyzed offline using ChartPro (AD Instruments).

Heart rhythm analysis on data from implanted transmitters was performed offline using Ponemah (Data Sciences International, St Paul, MN). 48 hours of heart rhythm data were analyzed for each mouse, with determination of average heart rate and PR interval. To generate heart rate histograms, two 12-hour segments corresponding to low activity (8AM-8PM) and two 12-hour segments corresponding to high activity (8PM-8AM) were analyzed for each mouse. The average heart rate for each minute over the 48-hour period was sorted into 10-bpm bins, with the data from low and high activity phases plotted and analyzed separately. To measure the intrinsic heart rate, a 5-minute period of stable heart rate after injection of atropine and propranolol was averaged for each mouse tested. Similarly, maximum heart was determined by averaging over 1 minute of continuous and stable heart rhythm after injection of 1.0 μg isoproterenol.

For identification of arrhythmias, RR intervals that exceeded twice the RR average of the low- or high-activity period in which they occurred were identified automatically using Ponemah. If an RR interval was longer than twice the average of the previous three beats, it was classified as a sinus pause or, if a P wave was present without a QRS complex, an episode of AV block. Flagged RR intervals that did not meet the criteria for a pause but that exceeded twice the average RR were classified as bradycardic episodes. Low activity and high activity periods for each mouse had separate RR cutoffs determined by the average RR interval from 8AM-8PM (low activity) and 8PM-8AM (high activity). Animals of each genotype were grouped and between group statistical comparisons were made with the Mann-Whitney test with $p < 0.05$ was deemed significant.

Heart rate variability analysis was performed using Ponemah and MATLAB as previously described with minor modifications [10]. Low and high activity periods were analyzed separately. Poincare plots for a representative WT and *Cckar*$^{-/-}$ mouse were created using 2 minutes of consecutive heart beats acquired during an interval of stable rhythm in a low activity period. For time-domain analysis, three 10-minute segments were selected from a 2-hour time window for each mouse (12PM -2PM for low activity period and 12AM – 2AM for high activity period). Outliers greater than 2 standard deviations from the mean were removed. The average NN interval, standard deviation of all RR intervals (SDNN), root mean square of the difference of successive RR intervals (RMSSD), and percentage of normal consecutive RR intervals that differ by more than 6 msec (pNN6) were calculated in Ponemah. Statistical comparisons were made with the Mann-Whitney test and $p < 0.05$ was deemed significant.

##### *Isolation of Adult Mouse Cardiac Pacemaker Cells*

Excised hearts were placed in normal Tyrode's solution, which contained (in mmol/L): 140 NaCl, 5.4KCl, 1.0 $CaCl_2$, 1.2 $KH_2PO_4$, 5.0 HEPES, 1.0 MaCl2, 5.55 glucose, pH7.4 with NaOH. The SAN was isolated by excising the region of the right atrium bounded by the superior vena cava, inferior vena cava, interatrial septum, and crista terminalis. Isolated SAN tissue was cut into 1 mm wide strips and placed in the Ca2+ free Tyrode's solution with liberase (0.18 mg/ml, Roche, CA, USA) and elastase (2.0 U/ml) at 37 degree for 25 min. The SAN tissue was further dissociated to a single cell suspension by trituration with a glass pipette in KB solution, which contained (in mmol/L): 25 KCl, 10 $KH_2PO_4$, 5 HEPES, 20 Glucose, 20 taurine, 100 K-glutamate, 10 K-aspartate, 2.0 $Mg_2SO_4$, 5.0 Creatine, 0.5 EGTA, pH7.2 with KOH. The cell suspension was stored at 4°C, and re-equilibrated for at least 30 minutes at room temperature immediately before electrophysiological recording.

##### *Whole Cell Patch Clamp*

Action potentials were acquired from spontaneously beating adult cells of the indicated genotypes via the perforated patch-clamp technique with an Axopatch-700B amplifier at 32°C or 35°C using pCLAMP10.3 (Molecular Devices/Axon Instruments, Sunnyvale, CA) for data acquisition. All recordings were digitized at 20 KHz and filtered at 1.0 kHz. Electrodes had resistances of 3.5-4.0 MΩ. The bath solution was normal Tyrode's solution. The pipette solution contained (in mmol/L): 130 KCl, 10 NaCl, 0.04$CaCl_2$, 2 $Mg_2$ATP, 7 phosphocreatine, 0.1NaGTP, 10 HEPES and 200ug/ml of Amphotericin B (pH 7.2 with KOH). The pCLAMP 10.6 Clampfit module was used for analysis of action potential parameters. Firing rate was averaged from a 10 sec gap-free interval of stable recording. The change in firing rate was continuously recorded before and after the application of sCCK-8 (TOCRIS, cat# 1166) diluted in Tyrode solution at indicated concentrations.

The pCLAMP 10.6 Clampfit module and LabChart7 were used for analysis of action potentials (APs). To obtain the early diastolic depolarization velocity (EDDR), a take-off potential (TOP) was defined as the membrane potential when the dV/dT reached 10% of the maximum of the first derivative of the AP tracing ($dV/dt_{max}$). The total diastolic duration was defined as the time between the minimum diastolic potential (MDP) and TOP. Early diastolic depolarization (EDD) was defined as the time between 10% and 50% of the total diastolic duration. The EDDR was then defined as the change in membrane voltage in early diastole divided by the duration of EDD. Statistical comparisons of AP parameters before and after application of s-CCK8 were made with paired t-tests after each distribution passed a test of normality. $P < 0.05$ was deemed statistically significant.
